# Algorithmic Reconstruction of GBM Network Complexity

**DOI:** 10.1101/2021.09.21.461255

**Authors:** Abicumaran Uthamacumaran, Morgan Craig

## Abstract

Glioblastoma (GBM) is a complex disease that is difficult to treat. Establishing the complex genetic interactions regulating cell fate decisions in GBM can help to shed light on disease aggressivity and improved treatments. Networks and data science offer alternative approaches to classical bioinformatics pipelines to study gene expression patterns from single-cell datasets, helping to distinguish genes associated with control of differentiation and thus aggressivity. Here, we applied a host of data theoretic techniques, including clustering algorithms, Waddington landscape reconstruction, trajectory inference algorithms, and network approaches, to compare gene expression patterns between pediatric and adult GBM, and those of adult glioma-derived stem cells (GSCs) to identify the key molecular regulators of the complex networks driving GBM/GSC and predict their cell fate dynamics. Using these tools, we identified critical genes and transcription factors coordinating cell state transitions from stem-like to mature GBM phenotypes, including eight transcription factors (OLIG1/2, TAZ, GATA2, FOXG1, SOX6, SATB2, YY1) and four signaling genes (ATL3, MTSS1, EMP1, and TPT1) as clinically targetable novel putative function interactions differentiating pediatric and adult GBMs from adult GSCs. Our study provides strong evidence of the applicability of complex systems approaches for reverse-engineering gene networks from patient-derived single-cell datasets and inferring their complex dynamics, bolstering the search for new clinically relevant targets in GBM.

## 1. INTRODUCTION

Glioblastoma (GBM) is the most lethal pediatric and adult brain tumour. Despite advances in treatment, recurrence will occur in all GBM patients, and mean survival is only 15 months (Alifieris and Trafalis, 2015). GBM is a morbid disease that is driven by a high degree of heterogeneity and phenotypic plasticity in response to the interactions with their tumor microenvironment (Jung et al., 2019). The cell fate transitions and cellular decision-making in GBM cell populations are regulated by the dynamics of complex signaling networks (Suvà et al., 2014; Jia et al., 2017). Recent advances linking single-cell datasets and computational algorithms have improved our understanding of these complex networks and their orchestration of cell fate decisions of GBM transcriptional states (phenotypes) (Jin et al., 2018; Iacono et al., 2019). Despite this progress, quantitative approaches that reconstruct the information flow and dynamics of these complex networks remain under-applied. Pediatric GBM exhibits molecular patterns and collective behaviors which are fundamentally different from those of adult GBM (Paugh et al., 2010; Jones et al., 2017; Schwartzentruber et al., 2012; Sturm et al., 2012). There is a greater epigenetic burden in pediatric GBM marked by specific histone H3.3 modifications and aberrant DNA methylation profiles (Scwartzentruber et al., 2012; Sturm et al., 2012; Lulla et al., 2016; Harutyunyan et al., 2019). However, the complex signaling dynamics distinguishing pediatric and adult GBM subgroups, and the similarities within the molecular networks driving their cancer stemness, remain poorly investigated (Paugh et al., 2010; Jones et al., 2017). Answering the question of whether the reconfiguration of these underlying signaling networks in both GBM groups steers their cell fate dynamics would allow for the prediction of causal patterns in the disease progression and therapeutic responses.

Glioma-derived stem cells (GSCs) are believed to be a small subset of GBM cancer cells that largely contribute to emergent GBM adaptive behaviors such as phenotypic plasticity, clonal heterogeneity, self-renewal, aggressiveness (resilience), relapse/recurrence, and therapy resistance (Jung et al., 2019, Xiong et al., 2019). However, many different phenotypes in the tumor microenvironment, including immune cells, healthy cells, extracellular matrices, and blood vessels, form complex feedback loops with malignant GBM cells (Jung et al., 2019, Xiong et al., 2019). GSCs form complex networks with their tumor microenvironment. Signaling dynamics within this microenvironment and its reconfiguration govern the fitness and stemness of GSCs. A lack of quantitative understanding of the causal mechanisms (gene expression patterns) underlying GSC cell fate choices and transitions to their mature phenotypes hinders successful clinical interventions in the treatment of GBM (Jung et al., 2019, Xiong et al., 2019; Yabo et al., 2021).

Statistical approaches are traditionally used to study cell fate dynamics and infer complex networks from large-scale single cell transcriptomics by differential expression analysis through a combination of single cell data processing and clustering algorithms (Iacono et al., 2019).However, these algorithmic pipelines are inadequate for capturing the complex patterns and emergent behaviors of cancer network dynamics. Further, fundamental limitations associated with the raw counts of the scRNA-Seq complicate the inference of networks in complex diseases like GBM. These limitations include drop out events (zero counts), and the inherent noise and sparsity of single cell data. To extract quantitatively meaningful differences between GSC and GBM networks, while retaining the essential information representative of their complex dynamics, requires tools from the interdisciplinary paradigm of *complex systems theory*.

Complex systems theory, or complexity science, is the study of irreducible systems composed of many interacting parts in which the systems exhibit emergent behaviors. Emergence denotes systems in which the nonlinear interactions between the system and its environment give rise to complex patterns and unpredicted collective dynamics (Wolfram, 1988; Shalizi, 2006). Some general properties of complex systems include nonlinear dynamics, adaptive processes, self-organized structures, interconnectedness, collective behaviors, pattern formation, fractality, sudden phase-transitions, computational irreducibility, non-locality, long-term unpredictability, undecidability and multi-scaled, multi-nested feedback loops (Wolfram, 1988; Shalizi, 2006). The presence of multi-scaled feedback loops, in particular, is the defining feature of complex networks (Thurner et al., 2018). Traditional reductionist approaches are inadequate to quantify the properties and temporal behaviors of complex networks (Wolfram, 1988; Shalizi, 2006). Complex systems theory advocates the use of computational algorithms and tools from network science to dissect these complex networks (Thurner et al., 2018; Huang et al., 2009; Barabási and Oltvai, 2004).

The molecular networks coordinating the emergence of GSC and GBM phenotypes are such complex networks. To reveal the mechanisms underlying GSC cell fate decisions and transitions to their mature GBM phenotypes, we deployed several approaches from complex systems theory on data from single-cell RNA Sequencing (scRNA-Seq) count matrices. We compared pediatric GBM to adult GBM to identify the signaling network patterns distinguishing pediatric and adult GBM from GSCs. For this, we relied upon clustering algorithms, Waddington landscape reconstruction, multivariate information theory, network science (graph theory), and machine learning algorithms to map possible *cell fate dynamics* and identify robust expression markers (critical TFs and genes) driving the complex networks underlying GBM/GSC cell fate control and regulation. We found that distinct gene expression signatures regulate the cell fate decisions in the GBM and GSC patient groups we studied. In particular, we identified a set of key gene targets as master regulators of cell fate decision dynamics in all patient groups, and the critical drivers of GSC stemness networks. Mapping their energy landscape dynamics and cell fate trajectories in pseudotime (cellular transition dimension), we represented the GSC/GBM cell fate decisions as dynamical systems which allowed us to identify genes such as GATA2, FOXG1, SATB2, YY1, and SOX6, amidst others, as master regulators of information flow in their signaling networks. Our results help to understand how cellular fate decisions in GBM, identify potential drug targets for precision oncology, and provide a roadmap for data theoretic approaches to other such complex systems.

## 2. METHODS

### 2.1. General methodological framework

To understand GBM network complexity, we integrated several pediatric and adult IDH-wt GBM single-cell RNA-Seq (scRNA-Seq) datasets in an analytical pipeline that combines several network reconstruction and analysis tools (see subsections below). Details of the datasets used are provided in Table 1. Single-cell datasets were first filtered and normalized in a quality control step, and patient samples were removed from the scRNA-Seq counts expression matrix due to low unique molecular identifier (UMI)/high drop-out rates.

**TABLE 1.** Summary of single-cell datasets. The total number of patient samples (n) and number of single-cells within each patient group (N) used for each step of the clustering and single-cell trajectory inference process are shown.

| Patient Group | Single-Cell Dataset | # Patient Samples (n) and Single-Cells (N) for Seurat/BigScale | # Patient Samples (n) and single-cells (N) for scEpath Analysis | # of Cell Fate Trajectories in scEpath Waddington Landscape |
| --- | --- | --- | --- | --- |
| **Pediatric GBM** | Neftel et al. (18) | n = 7<br>N = 1850 | n = 7<br>N = 1850 | 2 |
| **Adult GBM** | Neftel et al.<br>(18) | n = 18<br>N ~ 21,500 | n = 7<br>N = 2221 | 4 |
| **Adult GSC** | Richards et<br>al. (19) | n = 28<br>N ~ 69, 000 | n= 13<br>N = 1504 | 2 |

Next, gene expression matrices were analyzed independently using the various clustering and trajectory inference algorithms discussed below. Here we provide a short summary (Figure 1). For the Seurat algorithm, the top 10 principal component analysis (PCA) loadings were used for the differential marker discovery; the top 25 PC loadings were used for the BigScale analysis. To identify the differential markers expressed in all clusters, the top 10 markers within these PC loadings were pooled and analyzed on the UMAP/tSNE patterning space of the cell fate clusters for each patient group. Similarly, the top 2 PCA loadings were used by the scEpath pseudotime analysis. The normalized scRNA-Seq counts of the discovered markers from the Seurat and BigSCale algorithms were pooled together, and separately analyzed for each patient group. The expression counts of these markers were then run through the PIDC Network Inference algorithm to obtain gene receptor networks.The differential transcription factors identified in the pseudotemporal progression heatmaps were selected for scEpath analysis. Only the markers specific to each patient group were selected for the PIDC network inference. Lastly, complex networks analysis was performed on the reconstructed networks using transitivity and centrality scores to assess the network structure and dynamics (information flow) to identify key regulators of GBM/GSC cell fate decisions. Further, algorithmic complexity measures, as provided in the Supplementary Material, were used to identify gene markers which could accurately discriminate the patient groups by machine learning classifiers. Within the established gene networks, algorithmic complexity was used to identify robust discriminants that could accurately distinguish the three patient groups (i.e., pediatric GBM, adult GBM, and adult GSC), based on the performance of machine learning classifiers on their algorithmic complexity scores (see Supplementary Information).

**FIGURE 1.**
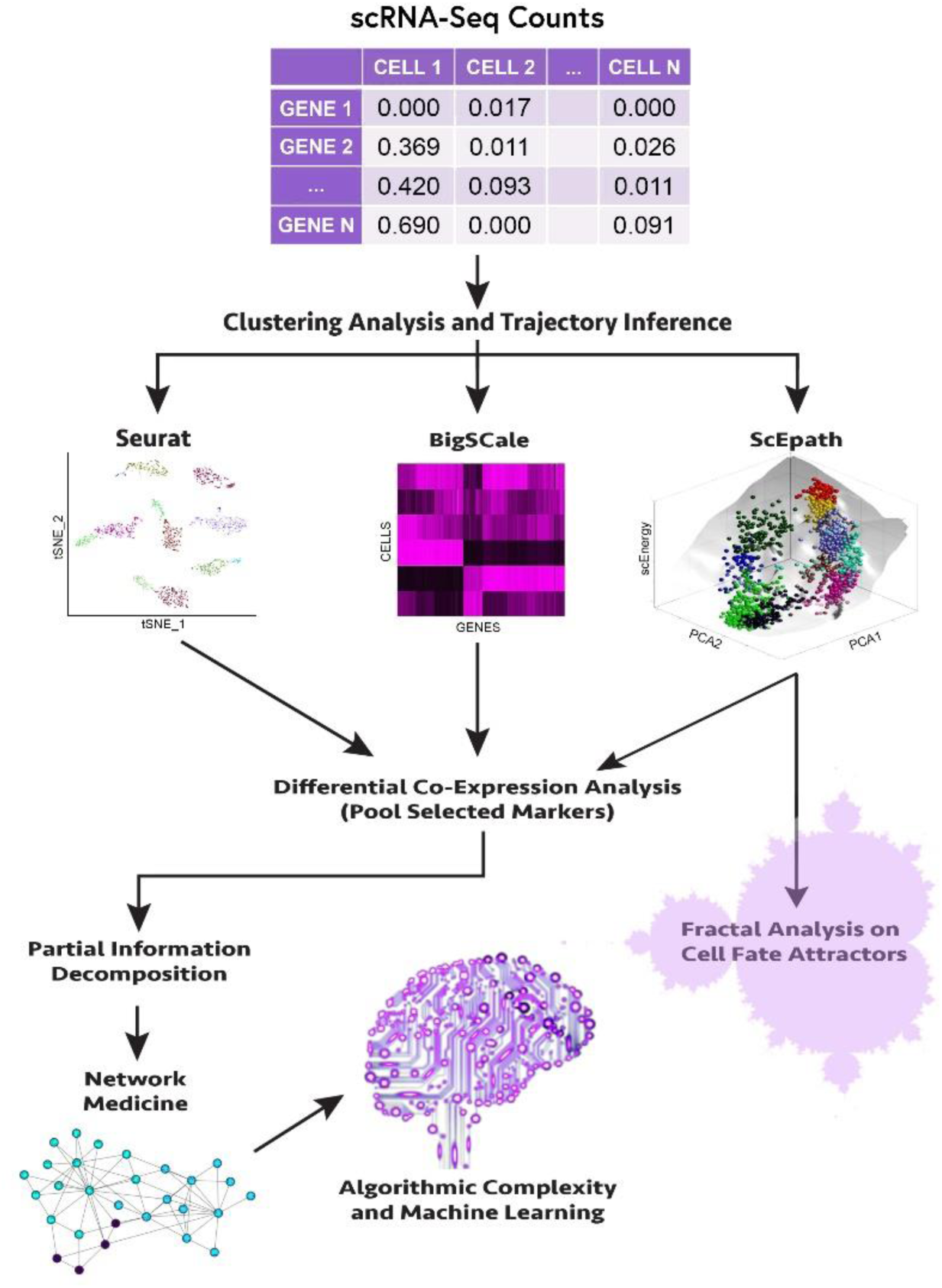
Workflow schematic of gene expression network pattern discovery. Flowchart summarizing the methodological approach to differential marker discovery and cell fate dynamics inference (see Methods section 2.1).

### 2.2. Single-cell datasets

Gene expression matrices for pediatric GBM, adult GBM, and adult GSC were obtained from the SingleCell Portal repositories from Neftel et al., 2019 and Richards et al., 2021 (Table 1). Briefly, GBM patient samples from Neftel et al. (2019) contained the single cell RNA-Seq counts of four phenotypes (or cellular states): macrophages, malignant GBM cells, oligodendrocytes, and T-cells. Adult GSC consisted only of stem cells. Overall, our dataset included 28 adult GSC datasets, 7 pediatric GBM, and 18 adult GBM scRNA-Seq expression count matrices.

As a quality control measure for the Seurat and BigSCale clustering, two adult GBM samples and one pediatric GBM sample were dropped in the filtering process (prior to clustering) due to high zero-counts (i.e., low UMI). Importantly, we confirmed that our findings were insensitive to the number of patient samples within each patient group: including these removed samples did not change the differential expression analysis. To further validate this finding, one sample was randomly chosen and dropped from the total number of samples from each patient group to verify whether the clustering analysis changed (i.e., leave-out-one cross-validation) and we confirmed the clustering results were identical. Beyond 2500 cells, the computational time complexity of the scEpath algorithm increased. Thus, the total cell counts of all three patient groups were kept at the maximum computational threshold for the scEpath analysis (see Section 2.3.3). Further, to visualize the cell fate attractor dynamics at the same fine-scale resolution for all patient groups, cell counts were kept roughly the same for each GBM type. Selecting a different combination of adult GSC samples did not change the scEpath landscape or results, as the trial of multiple random selections (> 6 distinct combinations) reproduced identical results. A complete description of the experimental approaches used to derive these datasets from their original studies is provided in the Supplementary Information.

### 2.3. Clustering techniques

Clustering algorithms were used to identify differential markers co-expressed within all patient groups and distinguish a robust network regulating the cell fate dynamics across all phenotypes.

#### 2.3.1. Seurat algorithm

scRNA-Seq count matrices were pre-processed to obtain normalized and binarized count expressions. Seurat initially performs a cluster analysis by principal component analysis (PCA) dimensionality reduction followed by a graph-based clustering (k-nearest neighbor (kNN) graph) based on the Euclidean distance of the 10 PCA loadings using the FindNeighbors function and Louvain community detection algorithm (modularity optimization) using the FindClusters function (parameter can be tuned between 0.4-1.2 for optimal results), to cluster cells by their Jaccard index-expression similarity (see Seurat Clustering tutorial in GitHub code). All clustering parameters were kept to their default settings. Next, the cells within the graph-based clusters were visualized on Uniform Manifold Approximation and Projection (UMAP) or t-Distributed Stochastic Neighbor Embedding (TSNE) space (i.e., unsupervised nonlinear dimensionality reduction techniques) (Stuart et al., 2019). Differential markers from the top 10 PCA loadings were visualized in UMAP space (analysis does not vary for TSNE space) using the FindAllMarkers function with parameters: min.pct = 0.25 and logfc.threshold = 0.25. We clustered similarly expressed cells together in the low dimensional space by finding differentially expressed features/markers corresponding to the highest ten PCA loadings in the graph-based clusters. To identify markers that govern disease progression and transcriptional dynamics, we imposed the condition that selected markers for the network reconstruction must be expressed in all clusters of the three patient groups (pediatric GBM, adult GBM, and adult GSC).

#### 2.3.2. BigSCale algorithm

BigSCale is a framework for clustering, phenotyping, pseudotiming, and inferring gene regulatory and protein-protein interaction networks from single-cell data (Iacono et al., 2019). A SingleCellExperiment class was created from the scRNA-Seq raw count matrices for BigSCale processing, and counts were replaced by z-scores. Cellular clustering was established by first computing all pairwise cell distances using the Pearson correlation to generate a distance matrix. Following, cells were assigned to cluster groups via the Ward’s linkage/method (an agglomerative hierarchical clustering algorithm). Iterative differential expression analysis was performed between the clusters of cells and the differential markers within the identified clusters were assessed using the getMarkers function (see BigSCale 2 tutorial in Github code). The markers specific to a cluster were sorted from the highest (most significant) to the lowest (least significant) z-score for the selection of cluster-specific differential and co-expressed gene markers within the top 25 PCA components. A z-score threshold of 5.0 was used as a cut-off threshold while the min_ODscore parameter was kept default at 2.33. This imposed cut-off acts as a filtering mechanism to retain only the markers with significant expression changes per cluster. As in the Seurat analysis, we imposed the condition that selected markers for the network reconstruction must be expressed in all clusters of the three patient groups.

#### 2.3.3. ScEpath algorithm

We applied single cell Energy path (scEpath) to reconstruct the 3D-energy landscape of cells and infer regulatory relationships from their transcriptional dynamics (Jin et al., 2018). scEpath is a Waddington Landscape reconstruction algorithm with an unsupervised clustering framework for cell lineage hierarchy mapping and studying the pseudotemporal transcriptional dynamics in cell fate decisions. In this trajectory inference algorithm, information flow and network reconfiguration underlying the cellular decision-making steer the topography of cell populations’ energy landscapes (also referred to as a cell fate landscape, attractor landscape, or Waddington’s epigenetic landscape (Waddington, 1957)). A cell state (cell fate) corresponds to a specific transcriptional (gene expression) program and phenotype of a given cellular population. Cell clusters higher on the energy landscape correspond to stem-cell like states (unstable attractors) with higher differentiation potency, while cell states stuck in lower energies (valleys, or stable attractors) correspond to differentiated (mature) phenotypes with lower potency/plasticity (Figure 2).

**FIGURE 2.**
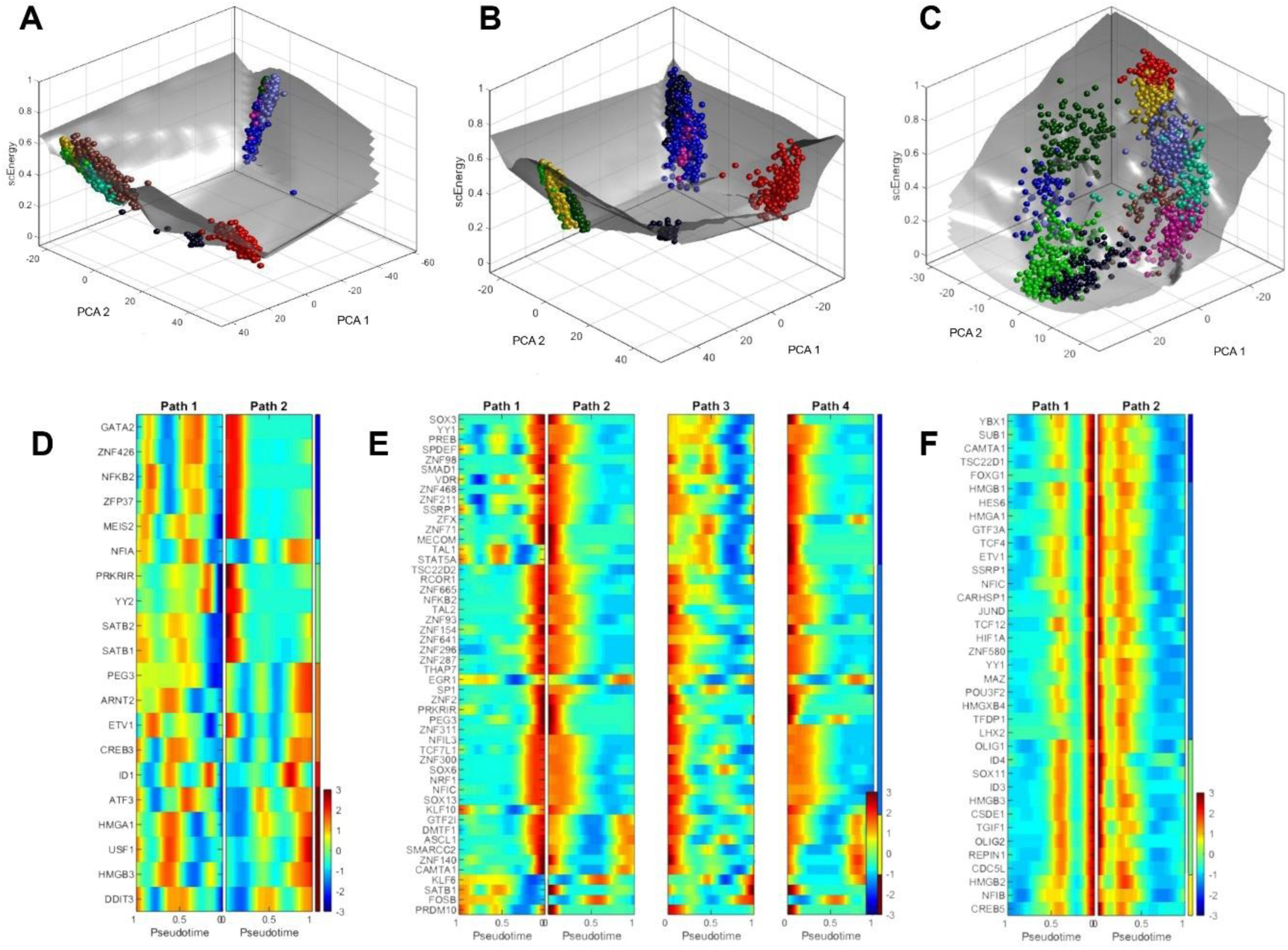
Waddington landscape reconstruction differentiates adult and pediatric GBM, and adult GSC critical genes and transcription factors for cell fate transitions. A-C) The distinct phenotypes of each patient group were clustered on the Waddington energy landscape by their similarity in gene expression. The cell clustering patterns on the landscape are referred to as *attractors*. Balls represent the transcriptional states (cell fates) and paths correspond to the cell fate differentiation trajectories on the Waddington landscape. A) Pediatric GBM. B) Adult GBM. C) Adult GSC. D-F) Heat maps of the critical transcription factors involved in the differentiation and cell fate transitions between the distinct attractors (phenotype clusters) of the GBM/GSC Waddington landscape. The color gradient represents the intensity of the gene expression in pseudotime trajectory, where blue implies low expression and red implies high expression of the gene (TF) during the cell fate choices along the cell differentiation trajectories. The path corresponds to the inferred trajectories in between the cell state attractors on the Waddington landscape. D) Pediatric GBM. E) Adult GBM. and F) Adult GSC.

scEpath allows for the visualization of cell fate transition probabilities in the population, mapping of cell lineage trajectories in pseudotemporal ordering, and inference of cell fate decisions from patient-derived scRNA-Seq datasets using the following steps: (i) preprocessing of scRNA-seq count matrix, (ii) gene regulatory network (GRN) inference, (iii) single cell energy (scEnergy) calculation, (iv) 3D energy landscape reconstruction via principal component analysis and structural clustering; (v) Transition probabilities calculation, (vi) Inference of cell lineage hierarchy via a probabilistic directed graph, (vii) pseudotime trajectory inference and, (viii) downstream analyses of identifying critical transcription factors (TFs) governing the cell-fate commitments (Jin et al., 2018). A detailed description of the scEpath algorithm is provided in the Supplementary Information.

To perform the scEpath analysis on our data, we first pre-processed the log-normalized (within patient-groups) count matrices with respect to their gene expression values by filtering out zero counts. The differential markers were selected from the first two significant PCA components. We then ran the scEpath MATLAB code from (Jin et al, 2018) on these processed datasets. GSC patient samples BT127, BT48, and BT84 from Richards et al. (2021) were used for all scEpath analyses on GSC. Seven pediatric GBM samples from (Neftel et al., 2019) and seven adult GBM samples, selected to match the cell count of the pediatric patient group, from (Neftel et al., 2019), were analyzed. We confirmed that the number of patients did not influence the results and analysis by selecting different random sets of adult GBM samples. We then ran energy (Waddington) landscapes reconstruction on the following population sizes: pediatric GBM: n= 7, N= 1850 cells; adult GBM: n = 7, N= 2221 cells; adult GSC: n=3, N=1504 cells.

scEpath smooths the average normalized expression of each gene using cubic regression splines to map the pseudotemporal gene expression dynamics along the inferred trajectories of the cell fates on the landscape, leading to smoothed gene expression along a lineage path (Jin et al., 2018). Leveraging this, we inferred key regulatory TFs for the cell fate differentiation by considering all PDG genes with a standard deviation > 0.5 and a Bonferroni-corrected p-value below a significance level α = 0.01 for the expression greater than a threshold (e.g., log2(fold-change) > 1). The probabilistic-directed graph network and the cell lineage hierarchy inference parameters were kept at default settings (quick_construct = 1; tau = 0.4; alpha = 0.01; theta1 = 0.8). The pseudotime-dependent genes were identified using parameters sd_thresh = 0.5; sig_thresh = 0.01; nboot (see hyperparameter-optimized code in GitHub link).

#### 2.3.4. Fractal and multifractal analysis

We applied fractal analysis to quantify the complexity of the phenotypic patterns on the scEpath cell fate attractor landscape. Fractals are signatures of complex systems (Mandelbrot, 1982), and the fractal dimension is a non-integer, fractional dimension characterizing the statistical self-similarity and roughness of a pattern. A higher fractality in tumor structures may imply that the tumor is more complex, resilient (i.e., withstands environmental perturbations), aggressive, and difficult to treat (Coffey, 1998; Baish and Jain, 1998). As such, the fractal index provides a quantitative measure of the cell fates’ phenotypic plasticity (i.e., higher for stem cell-like fates) and disease progression.

We used ImageJ plugin FracLac (v2.5) to compute the fractal dimension (FD) of analyzed samples using the BoxCount algorithm on the cell state attractors (patterns of cellular distributions on the scEpath energy landscapes). To calculate the fractal dimension, landscape images were converted to black and white. Attractor fractal dimensions reconstructed from the cell fate landscapes found to be non-integer were considered to exhibit a fractional dimension in phase-space. Higher fractal indices indicate more complex dynamics that are irregular and asymptotically unpredictable, since in dynamical systems theory, patterns of systems exhibiting deterministic chaos have a fractal dimension (i.e., strange attractors) (Strogatz, 2015).

#### 2.3.5. Partial Information Decomposition and Context network inference

Using the differential expression markers identified by the various approaches discussed above, we reconstructed the underlying complex networks driving the GBM/GSC cell state dynamics on the Waddington energy landscapes. Network inference tools study the statistical dependencies between genes amidst distributions of expression levels in populations of sampled cells (Chan et al., 2017) by inferring a graph-theoretic representation of the functional relationships between the drivers of complex behaviors such as cell fate transitions, thus allowing for the quantification of the relationships between identified differential transition markers and tracking how these relationships change across distinct phenotypes. Partial Information Decomposition and Context (PIDC) networks have been suggested to outperform traditional gene regulatory network inference approaches using correlation metrics, mutual information, Boolean networks, or Bayesian inference methods for network reconstruction (Chan et al., 2017). We used this PIDC network inference algorithm to obtain a network structure of GBM and GSC samples.

The Julia packages *InformationMeasures*.*jl* and *NetworkInference*.*jl* were used to reconstruct the GRN networks. PIDC network inference uses partial information decomposition (PID) to infer regulatory interaction networks from gene expression datasets. We used the *NetworkInference*.*jl* package to establish the (undirected) networks from the multivariate information measure (PID) calculated from the gene expression matrices. Gene expression counts were first discretized via Bayesian blocks discretization and the maximum likelihood estimator (Chan et al., 2017). The PIDC network pattern is the simplest network the algorithm can construct such that the distance between the nodes (genes or TFs) are minimized given their weights (PID score). Network measures characterizing the structure, properties, and information flow of these complex networks were then computed and the most differentially expressed genes were identified by the clustering algorithms using PID scores.

#### 2.3.6. Block Decomposition Method Calculations

We evaluated the algorithmic complexity of key nodes (genes) of the inferred signaling networks to further identify robust markers distinguishing GBM and GSC. Algorithmic complexity is a complementary measure that identifies the minimal amount or set of information in our inferred complex networks which regulate the phenotypic plasticity dynamics across the patient groups, and as such the genes/TFs with highest algorithmic complexity could be robust disease screening tools in precision oncology. The K-complexity of a string *s, K*(*s*), also known as Kolmogorov or algorithmic complexity, is the shortest computer program length needed to output that string. This can also alternatively be interpreted as the length of the shortest description of a system (Zenil et al., 2016). Since *K*(*s*) does not depend on a choice of probability distribution like Shannon entropy, it is more robust for the assessment of system complexity (Zenil et al., 2016, Zenil et al., 2019). Formally, the Kolmogorov complexity of a discrete dynamical system is given by

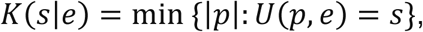

for a string or array *s*, where *p* is the program that produces *s* and halts running on a universal Turing machine *U* with input *e*. Then, K(s) is a function that takes a string or matrix *s* to be the length of the shortest program *p* that generates s. However, *K*(*s*) *is* in principle incomputable and must be approximated using the coding theorem method (Zenil et al., 2019). We therefore used the Block Decomposition Method (BDM) to approximate the *K*(*s*) of a dataset, which provides local estimates of the algorithmic complexity (Zenil et al., 2016). BDM is available in the online algorithmic complexity calculator [OACC] and its R-implementation (see Availability of Data and Material). The BDM is defined as

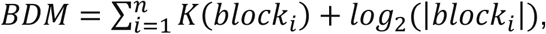

where the block size must be specified for the n-number of blocks. When the block sizes are higher, better approximations of the K-complexity are obtained (Zenil et al., 2016, Zenil et al., 2019).

To calculate the BDM, we selected scRNA-Seq counts of seven randomly chosen patient samples from each of the three patient groups. String length was kept the same for all gene candidates from each sample. Accordingly, we chose the cell count expressions of 46 cells from each patient sample for this analysis. The R-implementation of the Online Algorithmic Complexity Calculator was used to compute the BDM estimates of K-complexity for each expression string. scNA-Seq counts of the top gene interactions with highest PID scores were selected from each network and binarized. We then performed BDM on these binarized strings using a block size of 12 and alphabet size of 2 bits to estimate the K-complexity (i.e., BDM score) (see Supplementary Information for BDM Results).

## 3. RESULTS

### 3.1. Key driver genes mediating the cell fate transition dynamics in GBM/GSC epigenetic landscapes are identified using the scEpath algorithm

The Waddington landscape reconstruction identified causal patterns (attractors) to which the distinct transcriptional states within each patient group cluster (Fig 2A-C). Distinct patient group clusters were determined by the scEpath algorithm (colored by similarity in gene expression (i.e., phenotypes) in Figure 2). Three and four meta-clusters were identified in the pediatric GBM (Fig 2A) and adult GBM (Fig 2B), respectively while sub-populations are observed within each meta-cluster indicating the presence of phenotypic heterogeneity and epigenetic plasticity. Many genes encoding transcription factors (TFs) were identified as the transition genes required for cells to transition from one attractor to another. We mapped the expressions of these transition genes across the inferred cell fate trajectories (Fig 2D-F) and found similarities in the gene expression signatures and similar oscillatory patterns in EMP1, MTSS1, PHGDH and OLIG1/2 (Fig 3). These markers were selected in the clustering and trajectory inference process as explained above. Their similarity was assessed by their expression variation along the cell fate trajectories in pseudotime (Figure 3). We also identified OLIG1/2 as critical transcription factors in the adult GSC phenotypic transitions (Fig 2F).

**FIGURE 3.**
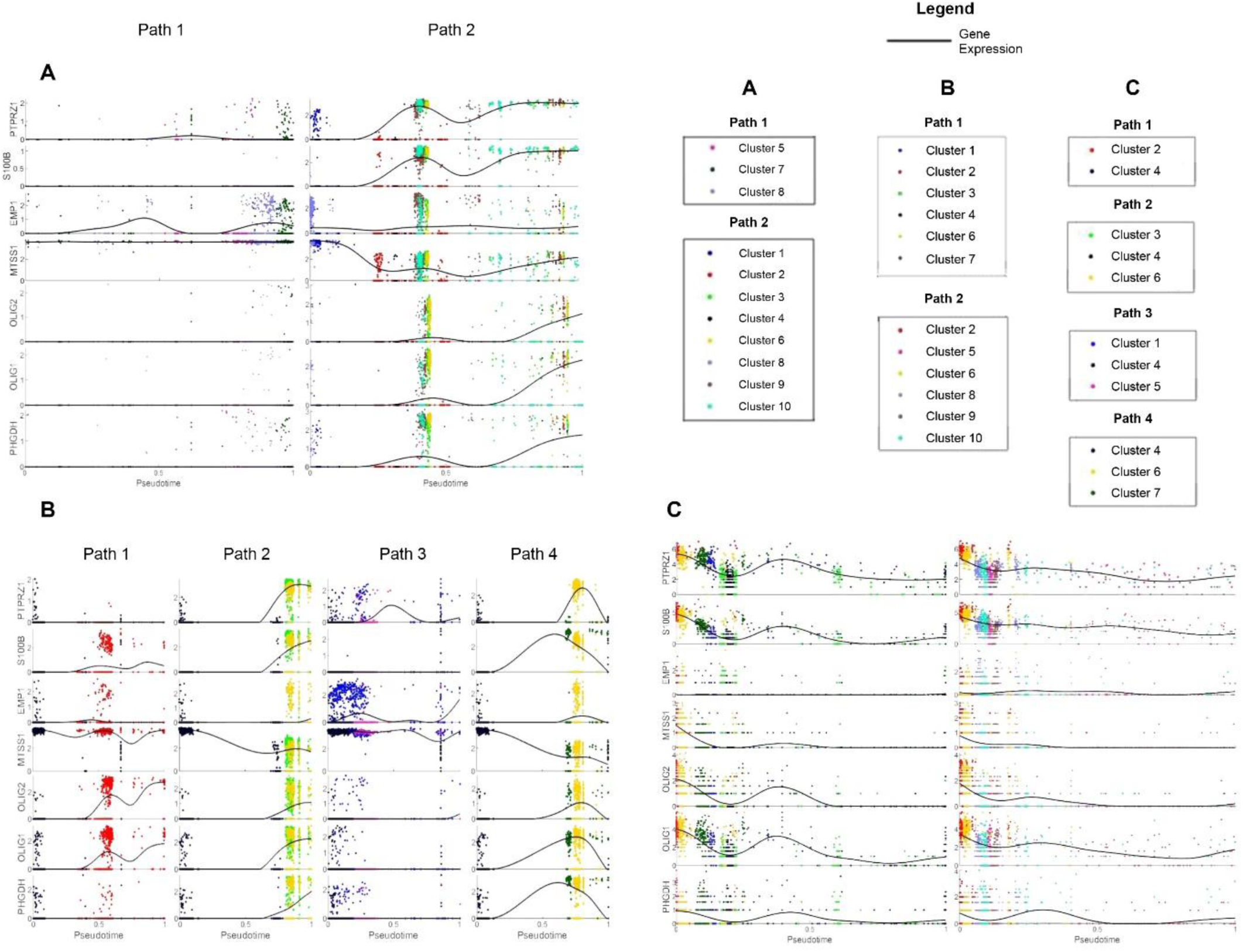
Reconstructing pseudotime dynamics in GBM/GSC cell fate decisions of the Waddington landscape. Average normalized gene expression in cells plotted along pseudotime after fitting with a cubic smoothing spline (black line). Cells are colored according to cell clusters defined by scEpath. The expression patterns of the top genes identified by scEpath and BigSCale algorithms (via correlation metrics) showed significant changes along the pseudotime trajectory inferred by scEpath algorithm. Selected gene markers in A) pediatric GBM, B) adult GBM, C) adult GSC.

### 3.2. Pseudotime expression dynamics identifies oscillatory patterns in critical gene targets

Given the key driver genes and transcription factors identified by scEpath trajectory inference, we next sought to infer similarities in gene expression dynamics during cell fate transitions within each patient group amidst the identified critical gene markers. Using clustering algorithms (see Methods), we found that PTPRZ1 and S100B showed nearly identical expression dynamics in pediatric GBM along both cell fate trajectories on the Waddington landscape, whereas genes such as EMP1, MTSS1, and PHGDH had more complex dynamics and exhibited oscillations during cell fate dynamics in pediatric GBM and adult GSC (Fig 3A and Fig 3C). The expression metric used to compare the dynamics of the different pseudotime-dependent genes correspond to the cubic spline smoothened average normalized expression along the pseudotime interval of [0,1].

In adult GBM, NACA and PABPC1, and TPT1 and PSAP had similar expression patterns across all four differentiation paths (Fig 3B). S100B, OLIG1, and PHGDH all had a broad expression profile in path 4 (Fig 3B). Furthermore, the presence of four cell clusters in adult GBM landscape (Fig 2B) is in good agreement with previous classifications of four molecular subtypes of adult GBM (Verhaak et al., 2010). The expression of EGFR and PDGFRA were distinctly higher in one of the four cell fate clusters/attractors (Figure S4B). However, the expression of IDH1 exhibited oscillatory dynamics in all four paths/attractors (data not shown). In adult GSC, many of the identified markers had similar gene expression profiles in pseudotemporal ordering (Fig 3C). For instance, PTPRZ1, NACA and PABPC1, were all found to have similar expression dynamics in both transition paths (Fig 3C). Notably, OLIG 1 and OLIG2 were found to have similar expression patterns in all three patient groups across all cell fate transition trajectories of the landscape (Fig 3A-C).

Notably, we identified that genes such as STMN3, MTSS1 and TAZ are critical regulators in one transition pathway, while PSAP, TPT1, and PTPRZ1 are relevant for the other transition trajectory on the pediatric GBM’s Waddington landscape (Fig 3A and S4A). The same trends in pseudotemporal gene expression patterns in STMN3 and PTPRZ1 have also been found in the adult GSC cell fate trajectories (see Fig S4C in the Supplementary Information). In all three patient groups, OLIG1, OLIG2, PHGDH, and TIMELESS had similar expression profiles within the distinct cell fate transition paths indicating potentially some network coordination or collective oscillations. Some signals (e.g., BCAN and CLU) were found to exhibit oscillations that may be indicative of complex dynamics with time-series expression analysis (Supplementary Information). These findings suggest that the identified markers involved in GBM/GSC cell fate decisions exhibit similar patterns in their expression dynamics, and that the identified critical genes are functionally putative master orchestrators of cell fate transitions/differentiation of the heterogeneous phenotypes within a GBM patient’s tumor.

### 3.3. PIDC Network Inference algorithm reconstructs the regulatory network configurations driving GBM/GSC cell fate transitions

We next reverse engineered the signaling networks coordinating the information flow in GBM and GSC using Partial Information Decomposition and Context (PIDC). Though the network topography may seem similar, the arrangement of the interactions from highest influence on the information flow (i.e., top PID scores) to those of the weakest interactions (lowest PID scores) vary for each patient group. As seen in Figure 4A, OLIG1 and OLIG2 have the highest PID score of 1.9508, followed by S100B and PTPRZ1 interaction with a PID score of 1.9303 in pediatric GBM, suggesting a strong relationship between these two genes in the complex network steering their cell fate decisions (Fig 4A). We found that S100A10 and EMP1 have the highest interaction in adult GBM with a PID score of 1.9517 (Fig 4B), whereas NACA and TPT1 had the highest interaction in adult GSC with a PID score of 1.9628 (Fig 4C). A distinct pattern was observed in the PIDC regulatory network of adult GSC sample BT127 (highest quality GSC cells). The highest interaction was observed between PHGDH and TIMELESS at a PID score of 2.762. Other top interactions identified for the TF networks (Fig 4E-G) had similar pseudotemporal expression dynamics (Figure S4 A-C in the Supplementary Information). ATF3 and DDIT3 were the top interaction markers from the critical TFs identified for pediatric GBM with a PID score of 1.971 (Fig 4E). EGR1 and FOSB in the adult GBM group (Fig 4F), and YBX1 and HMGB1 were identified as the top interaction TF markers, with PID score of 1.992 (Fig 4G). These results suggest the reconfiguration of the nodes within the same complex signaling network may characterize GSC cells from GBM cells and distinguish pediatric GBM from adult GBM cell fate dynamics.

**FIGURE 4.**
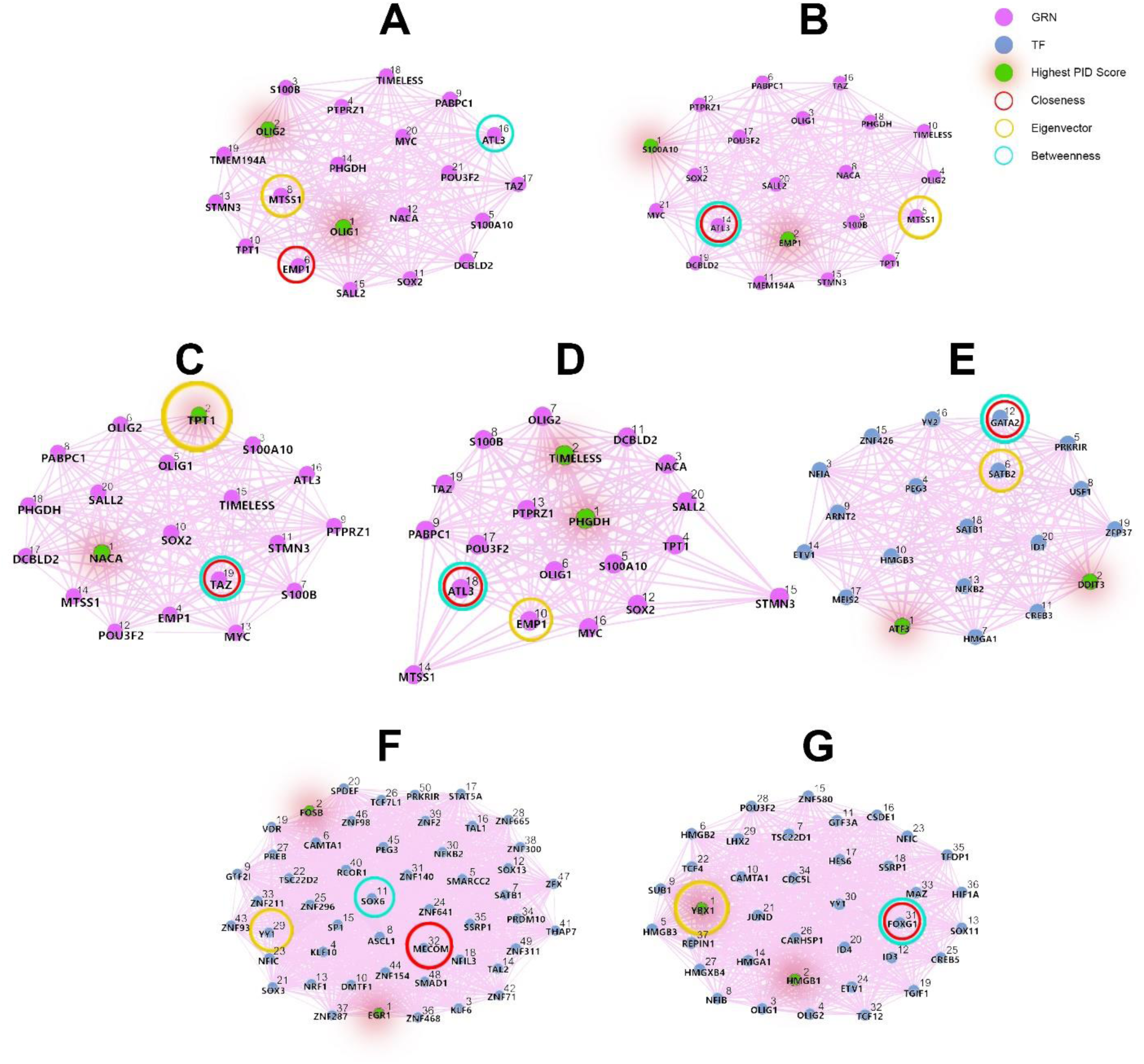
Mathematical modelling identifies key regulatory genes driving GBM networks. Gene regulatory networks of A) pediatric GBM, B) adult GBM, C) adult GSC, D) adult GSC sample BT127, E) pediatric GBM transcription factors, F) adult GBM transcription factors, and G) adult GSC transcription factors. In each, the signaling networks show the information flow between critical signals required for the complex cell fate dynamics. The GRN networks identified by Seurat and BigSCale are colored in violet nodes (Fig 4A-D) while the scEpath TF networks are colored in teal (Fig 4E-G). The ranks were assigned a priority index by the PID content as indicated by the numbers on the nodes. A high PID content implies a high mutual information (dependence) of those gene interactions in the information flow network. The number index on the nodes of the network correspond to the PID score in a decreasing order, where rank 1 denotes the top (highest) value. As shown in the legend, the nodes with the highest PID score are colored in green with a red shadow. Additionally, three different colored rings are used to identify the nodes of the networks with the highest network centrality measures as identified in Figure 5.

### 3.4. Network centrality measures identify master regulators of information flow across the regulatory networks underlying GBM/GSC cell fate decision-making

Centrality is a key property of complex networks that influences the network dynamics and information flow (Iacono et al., 2019). The nodes (genes or TFs) with the highest centrality in the regulatory networks are the most biologically important signals. By measuring network centrality, we identified the primary genes regulating communication flow across each of the pediatric and adult GBM, and adult GSC networks (Table 2). In particular, we calculated the global clustering coefficient that measures the total number of closed triangles (link density) in a network. A clustering coefficient at its maximal value of 1 indicates that the neighbors of the gene (node) *i* form a complete graph (i.e., they all connect to each other) versus the converse for a clustering coefficient of 0 (Barabási and Posfai, 2016). We observed a lower clustering coefficient of 0.94 for the BT127 network in Figure 4D. In the transcription factor networks reconstructed from the scEpath heatmaps (italic columns, Table 2), the GSC TF network had the highest diameter, while the GBM networks (both pediatric and adult) had smaller diameters. The diameter is relatively in the same order of magnitude for the PIDC networks reconstructed from the Seurat-BigSCale markers (bold columns, Table 2) as they correspond essentially to the same set of genes interactions. The degree of centrality of all networks in Figure 4 was 1.0 at all nodes, except for the BT127 PIDC network which had a degree centrality of value of 1.0 only at nodes 1, 5, 10, 12, 13, and 16, and a clustering coefficient of value 0.96. The degree centrality of nodes 2, 7, and 8 were 0.89, the degree centrality of nodes 14 and 15 were roughly 0.5, and the remaining nodes had a degree centrality of 0.95.

**TABLE 2.** General Properties of Inferred Complex Networks. *G*(*V, E*) denotes the graph with the number of vertices *V* (the genes) and number of edges *E* for each inferred GRN network. The center values designate to the node index (gene) acting as the center of the simple weighted network. The clustering coefficient captures the degree to which the neighbors of a given node link to each other.

| **Network Properties** | **Pediatric GBM** | **Adult GBM** | **Adult GSC** | **BT127 (Adult GSC)** | *Pediatric GBM (TF Network)* | *Adult GBM (TF Network)* | *Adult GSC (TF Network)* |
| --- | --- | --- | --- | --- | --- | --- | --- |
| $G(V, E)$ | (21,210) | (21,210) | (20,190) | (20,174) | (20,190) | (50, 1225) | (37,666) |
| Center | 16 | 14 | 19 | 18 | 8 | 15 | 31 |
| Diameter | 3.037 | 3.283 | 3.490 | 3.106 | 1.869 | 1.474 | 3.936 |
| Global Clustering Coefficient | 1.00 | 1.00 | 1.00 | 0.94 | 1.00 | 1.00 | 1.00 |

The closeness centrality identified genes/TFs occupying a central position in a network (Iacono et al., 2019). The nodes corresponding to the highest closeness centrality for each GRN network were found to be Node 6 (EMP1) for pediatric GBM, Node 14 (ATL3) for adult GBM, Node 18 (ATL3) for GSC BT127, and Node 19 (TAZ) for GSC with closeness values of 1.398, 1.361, 1.006, and 1.184, respectively (Fig 5A).

**FIGURE 5.**
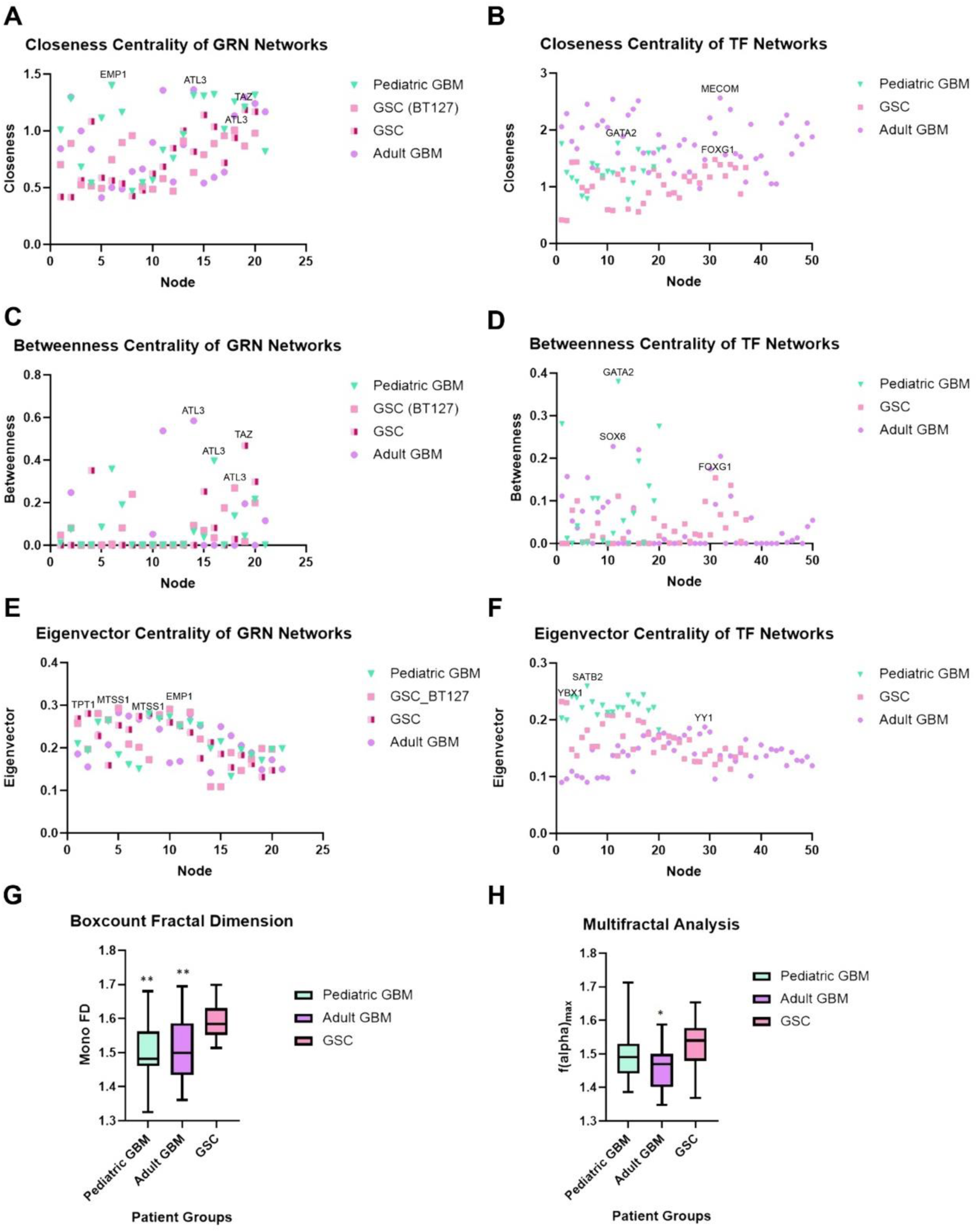
Centrality measures distinguish master regulators of information flow in GBM networks. *Three network centrality measures are assessed on the reconstructed GBM/GSC networks*. Three different network centralities were computed on the reconstructed networks: closeness, betweenness, and eigenvector centrality. The genes (nodes) occupying the highest of these centrality measures correspond to critical nodes steering the information flow in the complex signaling networks governing GBM/GSC cell fate transition dynamics. A) Closeness centrality of inferred GRNs. B) Closeness centrality of TF networks. C) Betweenness centrality of gene regulatory networks. D) Betweenness centrality of transcription factor networks. E) Eigenvector centrality of gene regulatory networks. F) Eigenvector centrality of transcription factor networks. G) Fractal dimension of cell state attractors on scEpath energy landscapes. A p-value of 0.0031 between the adult GSC and adult GBM, and p= 0.0011 between adult GSC and pediatric GBM was calculated for the box-count algorithm’s fractal dimension scores using the Kolmogorov-Smirnov test. Multifractal analysis of cell fate attractors on scEpath Waddington landscapes.

Nodes corresponding to the maximal closeness in the pediatric GBM, adult GBM, and adult GSC TF networks were found to be node 12 (GATA2), node 32 (MECOM), and node 31 (FOXG1), respectively with closeness measures of 1.761, 2.563, and 1.478 respectively (Fig 5B).

Betweenness centrality indicates the presence of regulatory bottlenecks (Iacono et al., 2019; Latora et al., 2017; Rodrigues, 2019). In our analyses, the highest betweenness measures for the pediatric GBM, adult GBM, BT127 adult GSC, and adult GSC GRN networks were node 16 (ATL3), node 14 (ATL3), node 18 (ATL3), and node 19 (TAZ), respectively with betweenness values of 0.3947, 0.5842, 0.2690, and 0.4678, respectively (Fig 5C). The trends in maximal betweenness values (Fig 5C) were in good agreement with the nodes contributing to the maximal closeness values discussed in Fig 5A, indicating that identified nodes are critical targets governing the information flow in these complex networks. The highest betweenness values for the TF networks were found to be to node 12 (GATA2) for pediatric GBM, node 11 (SOX6) for adult GBM, and node 31 (FOXG1) for adult GSC, with values of 0.3801, 0.2279, and 0.1539, respectively (Fig 5D). The highest values of eigenvector centrality, a measure of information flow across the network, for the GRNs were found to be node 8 (MTSS1) for pediatric GBM, node 5 (MTSS1) for adult GBM, node 10 (EMP1) for BT127, and node 2 (TPT1) for GSC, with measures of 0.2796, 0.2827, 0.2909, and 0.2805, respectively.

The maximal eigenvector is a measure of the hub-score, i.e., the highest authority node of hub networks (Latora et al., 2017; Rodrigues, 2019). The maximal eigenvector centrality of the TF networks was found to be node 6 (SATB2) for pediatric GBM, and node 29 (YY1) for adult GBM, and node 1 (YBX1) for GSC, with values of 0.2594, 0.1874, and 0.2322, respectively. SATB2 is a nuclear matrix-associated protein involved in chromatin remodelling and transcription regulation during neuronal differentiation (Gyorgy et al., 2008). Interestingly, all transition genes with high centrality measures identified in our network analyses, including EMP1, MTSS1, ATL3, and TPT1 have a TF-binding site for YY1 (Stelzer et al., 2016; GeneCards, 2021) (see Table 3).

**TABLE 3.** Interactions between transition genes and transcription factors identified in network analysis. Amidst the critical transition genes listed, the first four were identified as the central regulators of information flow across the GBM/GSC regulatory networks, while PTPRZ1 and S100B were other differential markers identified in our analyses. The list is not inclusive of all possible gene-TF interactions but restricted to the analysis of only the high importance (i.e., highest network centrality measures) scEpath TFs identified in our findings. The TF-gene interactions were identified using the GeneCards human gene database (GeneCards, 2021).

| TRANSITION GENES | TRANSCRIPTION FACTORS |
| --- | --- |
| ATL3 | YY1, FOSB, SOX6, GATA2, ATF3, EGR1, MYC |
| MTSS1 | YY1, ATF3, MYC |
| EMP1 | YY1, FOSB, GATA2, ATF3, MYC |
| TPT1 | YY1, ATF3, FOSB, SOX6, EGR1, OLIG1/2 |
| PTPRZ1 | YY1, YY2, EGR1, NANOG, POUF51 |
| S100B | YY1, GATA2, EGR1, SOX6, MYC |

We also performed fractal analysis on the attractors (cell clustering patterns) in the scEpath Waddington landscapes. The fractal dimension scores obtained on the cell state attractors on the energy landscape were compared across all groups (pediatric GBM (n=7), adult GBM (n = 18), and GSC (n=28)). The mean fractal dimension scores of the pediatric GBM, adult GBM, and adult GSC groups were 1.502 ± 0.099, 1.509 ± 0.091, and 1.588 ± 0.051, respectively (Figure 5G). The FD scores of the two GBM groups were nearly identical, while a statistically significant difference was observed from the GSC group. The multifractal spectrum *f*(*α*) was extracted from the multifractal spectra of the individual cancer samples energy landscape (n=54) (Fig 5H). Only the difference between GSC versus adult GBM was found to be statistically significant (p=0.0201) by a Kolmogorov-Smirnov test. The pediatric GBM, and adult GBM and GSC groups had a maximal multifractal spectrum *f*(α) value of 1.499 ± 0.092, 1.462 ± 0.066, and 1.521 ± 0.075, respectively.

## 4. DISCUSSION

Here we applied a collection of data theoretic and complexity science approaches to single cell RNA-seq data from pediatric and adult GBM, and adult GSCs to distinguish genes regulating communication within these cellular populations. Our findings demonstrate the application of these tools for deciphering GBM/GSC signaling networks to understand how network configuration orchestrates information flow and determines cell fate dynamics.

Multiple clustering algorithms were deployed to cross-validate their findings and ensure that the differential markers extracted for network analysis were robust, complementary, and of high importance in cell fate transition/differentiation mapping. There is a high degree of heterogeneity displayed by GBM stem cells. The complementarity of our results in our independent and orthogonal approaches are outlined in Table 3 by the associations identified between the transition genes and the scEpath TFs. Our approach using distinct clustering techniques and verifying their matching or complementary results was deployed to minimize the effects of expression heterogeneity and validate our findings (Krieger et al., 2020).

Using scEpath, we identified three and four meta-clusters in the pediatric GBM (Fig 2A) and adult GBM (Fig 2B), respectively, while sub-clusters within each meta-cluster indicated the presence of phenotypic heterogeneity and plasticity. However, the number of meta-clusters was ambiguous in the adult GSC landscape (Fig 2C), as shown by the continuous progression from the higher energy state clusters (stem-like fates) to the lower energy states indicating the potential presence of a complex attractor. An alternative measure to assess the significance of the scEpath clustering is the transition paths (cell fate trajectories). We predicted that the number of clusters identified in the pediatric GBM group corresponds to the neuronal, astrocytic-mesenchymal, and oligodendrocytic lineages, mirroring the healthy brain’s neurodevelopmental hierarchy (Jessa et al., 2019; Couturier et al., 2020). Similarly, the four clusters identified in the adult GBM group correspond to the four groups identified by Neftel et al. (2019), namely the OPC-like (oligodendrocytic progenitor cell), NPC-like (neuronal progenitor cell-like), AC-like (astrocytic cell-like), and MES-like (mesenchymal cell) lineages. Further, the infiltrated immune cells (i.e., T-cells and macrophages) grouped into the MES-like state (Neftel et al., 2019). Pediatric GBM cells showed less differentiation than the adult GBM samples, as indicated by the higher energy cell-states, suggesting a closer resemblance to the GSC sample. The two cell fate trajectories observed in the adult GSC sample may correspond to the transcriptional gradient of two cellular states observed in the original study by Richards et al. (Fig 2C), which were shown to mirror normal neurodevelopment and inflammatory wound responses (Richards et al., 2021).

The cell fate trajectories along the scEpath Waddington landscape (Figure 2A-C) were determined by the transition probabilities of the probabilistic directed graph reconstructed from the cell fate clusters, where the weighted edges of the networks correspond to the average normalized gene expression (see Supplementary Information for additional details). scEpath used the minimum directed spanning tree to find the maximum probability flow and minimal number of edges along the network, since cell fates transition to lower energy states during differentiation. The resulting tree approximates the cell state transition network and infers the observed developmental trajectories/lineage structures. The weighted edges of the cell state transition network were found to be proportional to the gene expression values seen in Figure 3, where the number of developmental trajectories inferred are indicated by the path numbers in Figure 2. Thus, two cell fate trajectories were detected in the pediatric GBM and adult GSC samples while four developmental trajectories were observed in adult GBM.

In pediatric GBM, the expression of transcription factors in pseudotime was shown to be highly nonlinear. Certain genes, including GATA2, were even found to be oscillatory in one trajectory while demonstrating an increasing or decreasing gradient of expression along the other cell fate trajectory. Likewise, patterns of other critical transition genes (TFs) were identified along the attractor dynamics between the distinct transcriptional states of adult GBM and adult GSC cells. Further, we found that genes such as EMP1, MTSS1, PTPRZ1 and S100B exhibited distinct gene expression oscillations in one differentiation trajectory (path) over the other(s) (Figure 3). These genes were also found to have TF-binding sites for the scEpath identified TFs with the highest network centrality measures in our downstream analysis (Figure 4). Together, these findings are indicative of a highly interconnected network of gene-TF interactions governing GBM/GSC cell fate decisions, and further suggest that the information flow across the inferred networks may steer cell fate decisions towards complex attractors on the GBM/GSC Waddington landscape.

Using network centrality measures, we identified OLIG1/2, TAZ, GATA2, FOXG1, SOX6, SATB2, YY1, and gene targets ATL3, MTSS1, EMP1, and TPT1 as critical genes governing the cell fate dynamics of GBM and GSC cells (Fig 5A-F). Many of these signals are neurodevelopmental transcription factors involved in healthy brain development, essential for conferring and maintaining cancer stem cells (GSCs). Maximal centrality scores indicated that they are key regulators of the network information flow in both GBM groups and GSCs. The functional significance of these transcription factors (see Supplementary Information) suggests their critical role in stem cell decision-making and differentiation dynamics. Our findings indicate that these genes may be strong candidates for therapeutic interventions points for the treatment of GBM. Other signaling interactions such as PTPRZ1 and S100B were identified in our analyses as potent clinically druggable targets in the treatment of GBM. Furthermore, we predicted that GATA2 and MTSS1 may provide a common ground for interlinking leukemogenesis, the complex signaling dynamics of leukemia/lymphoma affecting children, and pediatric glioma/GBM (Menendez-Gonzalez et al., 2019, Schemionek et al., 2015).

BDM was used to distinguish which of the differential markers can accurately classify/differentiate the three patient group samples (see Supplementary Information). We identified FOSB, HMGB1 and EGR1 as differential signatures which can accurately predict the patient groups in our single-cell analyses (see SI). The algorithmic complexity measured by the BDM allowed for the identification of critical network genes differentiating GBM and GSC phenotypes with the minimal information. The rationale for using gene/TF markers’ BDM as a phenotypic discriminant is that the algorithmic complexity denotes the shortest algorithm or minimal set of information within the complex networks inferred required to classify the distinct patient groups. As such, the identified genes/TFs may be useful biomarkers for prognostic screening and disease phenotyping in clinical medicine.

From the transcription factor (TF) networks identified by scEpath (Table 3), we distinguished some TFs to form interactions with some of the differential gene markers, suggesting cellular reprogramming targets for controlling GBM cell fate dynamics. Our study therefore quantifies how these markers’ expression vary in the cell fate transitions from stem-like to mature phenotypes. For a discussion on the biological significance of key genes and transcription factors identified in our analyses, see the Supplementary Results in the Supplementary Information.

The cell fate transition markers identified in our study, including PTPRZ1, EMP1, S100B, and MTSS1, are in good agreement with the findings from the original studies (SCP393 and SCP503). Although some of the signatures we identified overlap with the differential expression patterns of the original studies, they did not compare the co-expression of these markers between GSC and GBM. Markers differentiating distinct cellular states have been previously investigated (for instance, the original study by Neftel et al. identified copy number amplifications of the CDK4, EGFR, and PDGFRA loci and mutations of the NF1 locus, each favoring one of the four GBM phenotypes (Neftel et al., 2019)). Our study instead analyzed the expression patterns which fluctuate or form a differentiation gradient across the distinct cell states. Further, while previous studies have associated the differentiation markers of GBM progression identified here, our study demonstrates their novel integrated application to elucidate the roles of these network biomarkers in GBM cell fate decisions and differentiation dynamics. Indeed, while many of the identified genes or TFs have been previously studied in the context of neurodevelopmental regulation and glioma cell fate dynamics, most of those selected in our analyses are not yet documented in glioblastoma cell fate control. As such, we propose the identified interactions in Table 3 may provide clinically relevant GBM-specific precision therapeutics, and that our network analyses provide a quantitative tool to characterize which of the markers were of high importance (i.e., high centrality measures) in cell fate control, plasticity regulation, and transition dynamics. Future studies should exploit tools from algorithmic complexity theory including algorithmic network perturbation analysis (i.e., quantify the BDM changes across a network by node or link deletion) to better elucidate the inferred network dynamics in cancer cell fate control and regulation.

While previous GBM gene regulatory network inference methods vary from our approaches, our findings are consistent with their results. For example, Sun et al. found 15 hub genes in GBM-specific miRNA-TF networks, including PDGFRA and SOX11, and 6 hub TFs (including GATA1) as key regulators of GBM dynamics (Sun et al., 2012). In our study, we also identified PDGFRA and SOX11 as hub genes of the inferred GBM networks, and found that GATA2, an alternate isoform, overlapped with these findings. However, Sun et al. (2012) did not compare GBM of different age groups nor consider GBM-derived stem cells for reconstruct their differentiation networks. Similarly, a network inference study by Ping et al. (2015) revealed 17 hub genes in GBM networks, including EGFR and PDGFRA, as gene signatures of the proneural GBM subtype, both of which were identified in our analyses. In another study, GSEA and IPA-based gene enrichment pathway analysis discovered TAZ as a key regulator of GBM networks (Bozdag et al., 2014), which was also identified as a master regulator of GBM differentiation dynamics in our analyses.

Using multi-omic analyses, Suva et al. distinguished OLIG2, POU3F2 SALL2, and SOX2 as hub genes of GBM stemness networks critical for their tumor-propagation potential (Suvà et al., 2014). Our findings identified OLIG2 as a master control gene of GBM differentiation dynamics and established a connection between SOX2 expression and the critical hub gene FOXG1. Further, some epigenetic profiling studies have shown that aberrant histone modifications and methylation profiles are molecular signatures driving pediatric GBM and distinguishing them from their adult counterparts (Jones et al., 2017, Lulla et al., 2016, Sturm et al., 2012). Sturm et al. (2012) revealed that the TFs OLIG1, OLIG2 and FOXG1 are the master regulators of the hub gene networks driving these oncohistone pediatric GBM variants (i.e., K27M and G34V/R). Similar findings were recently reported by Wang et al. (2021), who identified the same set of TFs as critical drivers of pediatric high-grade gliomas’ epigenetic landscapes. We identified all three TFs reported by Sturm et al. and Wang et al. in our network approaches as critical regulators of GBM cell fate dynamics. Thus, our findings recapitulate the complex network dynamics driving the oncohistone variants of pediatric GBM and validate and extend previous findings.

It should be noted that there is a good deal of heterogeneity within the single-cell datasets across and with the patient groups. The original datasets contained 8 pediatric GBM samples, 20 adult GBM samples, and 28 adult GSC samples. For the initial clustering (i.e., differential discovery using Seurat and BigSCale), samples--two adult GBM and one pediatric GBM--with the highest drop-out rate (i.e., zero counts) were removed as a data filtering and quality control step prior to normalization. Subsequently, the number of adult GBM samples in the scEpath analysis was randomly selected to closely match the cell count numbers of the adult GSC patient groups. The down-sampling of GSC samples was necessary since scEpath analysis has a computational limitation on the number of samples which can be processed (roughly 2500 cells). As noted in the Methods, selecting a different combination of GSC samples did not change the results and including the removed samples did not change the differential marker discovery or expression analyses. Indeed, the global clustering patterns remained the same although there was greater dispersion in the local sub-clusters in the Seurat and BigSCale pattern space. However, including all n=8 pediatric GBM patient samples generated a shorter list of transition genes with abrupt transitions between the distinct phenotypes.

A limitation of our study is that we did not have access to pediatric GSC cells, given that adult GSC data have only recently been described (Richards et al., 2021). There may be other hidden causal interactions interconnecting the nodes of the complex networks we inferred that were not identified due to lack of data. Further, the lack of time-series scRNA-Seq counts is a barrier to understanding the complex dynamics of GBM/GSC networks. The pseudotemporal dynamics consist of inferred cell fate trajectories in a dimensionality-reduced data space (i.e., PCA space) by transcriptional similarity of cell fates. Ribosomal proteins and certain cytoskeletal markers (housekeeping genes) were also not pooled with the differential expression signatures for network inference (Figure S1).

This proof-of-concept study provides a comprehensive method to dissect the cybernetics of cancer cellular ecosystems and their cell fate dynamics. Current bioinformatic pipelines in cancer data science largely fail to reconcile the complex dynamics and temporal features of GBM transcriptional states, as they either take a reductionist approach to inferring gene expression patterns or rely on statistical correlation methods. In contrast, our framework provides a pipeline for causal pattern discovery and thereby allows the prediction/forecasting of how the differentially expressed transition genes control and regulate cell fate decision-making. Further, our approach allows for the mapping of these cancer cell fate behaviors to information flow across the inferred complex networks. Thus, these causal inference tools shed light on emergent behaviors in cell fate decisions such as transcriptional heterogeneity from a dynamical systems perspective. As such, we propose our methodological framework may provide a complementary and potentially more useful means to assess how the heterogeneous cancer phenotypes exhibit adaptive (emergent) behaviors and help forecast their dynamic response to drug/therapeutic perturbations at the level of molecular interactions.

## 6. CONCLUSION

This study demonstrates the use of complex systems approaches in deciphering the cybernetics of GBM/GSC networks, and shows how signaling dynamics differ between pediatric GBM, adult GBM, and adult GSC populations. By identifying transcription factors and genes, our combined approach serves as one part of the precision medicine toolbox for the treatment of GBM, suggesting both precision therapeutic targets and GBM reprogramming factors.

Prospective studies should explore the use of artificial neural networks, including Deep Learning algorithms, for single-cell transcriptomic analyses. Further, causal inference-based network inference methods such as Bayesian networks and algorithmic information dynamics should be investigated for GBM regulatory networks reconstruction. The epigenetic regulation of our identified transcriptional networks must be explored using high-throughput multi-omics datasets. Our network approaches should be extended to protein-protein interaction networks, epigenetic networks, and metabolic networks to investigate multi-omic levels of GBM heterogeneity, including oncohistone variants (i.e., K27M, K36M, G34V/R) and IDH1/2-mutants observed in pediatric gliomas (GBM).

## DECLARATIONS

### DECLARATIONS STATEMENT

The authors declare no competing interests.

### ETHICS APPROVAL AND CONSENT TO PARTICIPATE

Not Applicable

### AUTHOR CONTRIBUTIONS

AU performed the algorithms, wrote, and edited the manuscript. MC supervised, wrote, and edited the manuscript.

### DATA AND CODE AVAILABILITY

The datasets supporting the conclusions of this article are available in the Single Cell Portal repository:

https://singlecell.broadinstitute.org/single_cell/study/SCP393/single-cell-rna-seq-of-adult-and-pediatric-glioblastoma#study-summary

https://singlecell.broadinstitute.org/single_cell/study/SCP503/gradient-of-developmental-and-injury-reponse-transcriptional-states-define-functional-vulnerabilities-underpinning-glioblastoma-heterogeneity#study-download

All Codes and Algorithms used for the single-cell data analysis are available in the project GitHub page:

https://github.com/Abicumaran/GBM_Complexity_I

## Software/Algorithms

### Seurat

Project name: Seurat V3

Project home page: https://github.com/satijalab/seurat/

Archived version: 10.1016/j.cell.2019.05.031

Operating system(s): Platform independent

Programming language: R

Other requirements: Not Applicable

License: GNU Public License (GPL 3.0)

Any restrictions to use by non-academics: Not Applicable

### BigScale

Project name: BigScale V2

Project home page: https://github.com/iaconogi/BigSCale2

Archived version: 10.1186/s13059-019-1713-4

Operating system(s): Platform independent

Programming language: R

Other requirements: C++

License: Not Applicable

Any restrictions to use by non-academics: Not Applicable

### scEpath

Project name: single-cell Energy path (scEpath)

Project home page: https://github.com/sqjin/scEpath

Archived version: 10.1093/bioinformatics/bty058

Operating system(s): Platform independent

Programming language: MATLAB

Other requirements: C++

License: Not Applicable

Any restrictions to use by non-academics: Not Applicable

### OACC

Project name: Online Algorithmic Complexity Calculator V3

Project home page: https://github.com/algorithmicnaturelab/OACC

Archived version: 10.1016/j.isci.2019.07.043

Operating system(s): Platform independent

Programming language: R

Other requirements: Not Applicable

License: GNU Public License (GPL 3.0)

Any restrictions to use by non-academics: Not Applicable

### Network Inference

Project name: NetworkInference.jl and Partial Information Decomposition (PID)

Project home page: https://github.com/Tchanders/NetworkInference.jl

Archived version: 10.1016/j.cels.2017.08.014

Operating system(s): Platform independent

Programming language: Julia

Other requirements: Not Applicable

License: MIT “Expat” License

Any restrictions to use by non-academics: Not Applicable

### Julia LightGraphs

Project name: LightGraphs.jl V1.3

Project home page: https://github.com/JuliaGraphs/SimpleWeightedGraphs.jl

Archived version: Not Applicable

Operating system(s): Platform independent

Programming language: Julia

Other requirements: Jupyter Notebook and HTML

License: MIT “Expat” License

Any restrictions to use by non-academics: Not Applicable

### SciKit-learn

Project name: Scikit-learn

Project home page: https://scikit-learn.org/ or https://github.com/scikit-learn/scikit-learn

Archived version: http://jmlr.org/papers/v12/pedregosa11a.html

Operating system(s): Platform independent

Programming language: Python (≥ V3.7)

Other requirements: NumPy (≥1.14.6), SciPy (≥ 1.1.0), joblib (≥ 0.11), threadpoolctl (≥ 2.0.0), Google Colab or Jupyter Notebook

License: 3-Clause BSD license

Any restrictions to use by non-academics: Not Applicable

### FracLac

Project name: FracLac V2.5

Project home page: https://imagej.nih.gov/ij/plugins/fraclac

Archived version: Not Applicable

Operating system(s): Platform independent

Programming language: Java

Other requirements: Not Applicable

License: National Institute of Health (NIH) Public License

Any restrictions to use by non-academics: Not Applicable

