## Supplementary Material for "Algorithmic Reconstruction of GBM Network Complexity"

**SUPPLEMENTARY INFORMATION TO: ALGORITHMIC RECONSTRUCTION OF GBM NETWORK COMPLEXITY**

### SUPPLEMENTARY RESULTS

#### Clustering Algorithms Distinguish Differentially Expressed Markers in GBM/GSC Clusters.

The first ten PCA loadings of differential gene expression markers for each patient group were assessed using Seurat and BigSCale (see Methods, Main Text and Clustering Tutorials in Code Section/GitHub). We identified PHGDH, ATL3, XIST, RFX4, EMP1, DCBLD2, COL16A1, TP73, TMEM194A and TIM as having the highest z-score by PCA separated clusters using BigSCale. Other signatures such as EGFR and PDGFRA were observed in the top 2 PCA loadings of GBM samples in both Seurat and BigSCale clustering. However, they were not expressed as highly in all clusters and hence, only a few genes were found be relevant during filtering when imposing the condition that the gene marker must be expressed in all patient groups and all cell clusters in TSNE/UMAP pattern space (Figure S1). The zero-counts were filtered from the scRNA-Seq counts and samples were log-normalized within their respective patient groups.

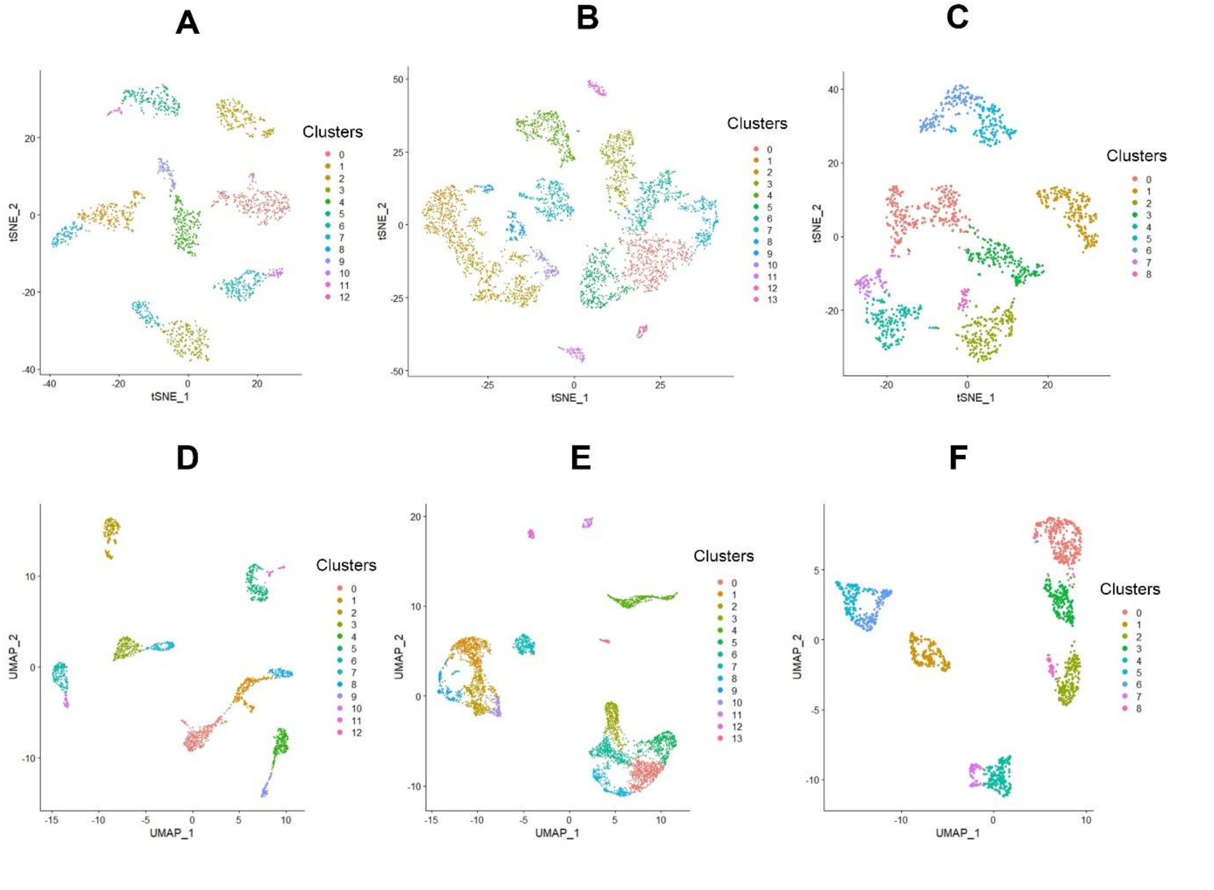


**Figure S1. Identifying differential expression markers from clustering patterns.** A) TSNE space of Pediatric GBM. B) TSNE embedding of Adult GBM. C) TSNE Space of Adult GSC. D) UMAP space of Pediatric GBM. E) UMAP space of Adult GBM. F) UMAP pattern of Adult GSC.

**Complex Cell fate Dynamics are inferred from Attractors on the Waddington landscape.**

Cell clusters were visualized on a two-dimensional contour plot as attractors of the Waddington landscape, the topology of which is defined as a function of their calculated scEnergy (Fig S2 A-C). We can see the light blue phenotypic cluster (*C8*), the global maximum on the pediatric GBM energy landscape, bifurcating towards two other global minima clusters (stable attractors) (Fig S2A). The complex signaling dynamics during the cell fate transition from the high energy cell-state cluster towards the two lower energy attractors indicate the possible presence of a complex global attractor, such as a strange attractor (Figure S4A), acting as a causal pattern connecting all clusters and driving GBM cell fate control and decisions. The same interpretation may apply to the adult GSC cells, where the red cell fate cluster (*C2*) was found to have the highest energy (Fig S2C). Their clustering patterns shows a bifurcation which seems to be interconnected by a global attractor. However, the adult GBM cells show four distinct phenotypic clusters, supporting the Verhaak classification of adult GBM molecular subtypes wherein the black cell cluster (*C4*) has the highest cell state energy (Verhaak et al., 2010). Our results thus suggest that pediatric GBM cell fate dynamics may more closely resemble adult GSCs than the mature GBM phenotypes in adults.

Cell lineage/state bifurcations were mapped as directed graphs by the scEpath algorithm (Fig S2 D-F). These bifurcations further suggest the complex dynamics underlying GBM/GSC cell fate decisions. The pseudotemporal ordering demonstrates an inferred temporal order based on similarities in their transcriptional states. Cell state transition probabilities to these possible phenotype clusters are shown as probabilistic directed graphs in Fig S2 D-F, where the numbers on the diagrams represent the transition probabilities from the high energy-state clusters to the low energy cell fate clusters. The high energy state clusters exhibit more stem-cell like behaviors while the low energy state clusters likely correspond to mature phenotypes. The inference of cell lineages were obtained by taking the Euclidean distance of pairs of cell cluster centroids as a weight of the edge. We found two cell fate trajectories in the pediatric GBM and adult GSC samples, and four in the adult GBM (Fig S2 D-F); refer to the Main Text for interpretation).

Lastly, the comparison of energy distributions among the identified cell clusters of each patient group are shown in FigS2 G-I. The significant reduction in scEnergy from the global maximum cell fate cluster to the local energy minima suggests developmental trajectories and cell fate differentiation/transition occurs from unstable high energy states to stable low-energy states (mature phenotypes). Without time-series datasets, it remains unelucidated whether a strange attractor(s) can interconnect the high and low energy states. Indicated p-values were calculated using a non-parametric two-sided Wilcoxon rank-sum test. The x-axes correspond to the distinct clusters visualized on the 3D energy landscapes in Figure 5 (Main Text and Figure S2 A-C).

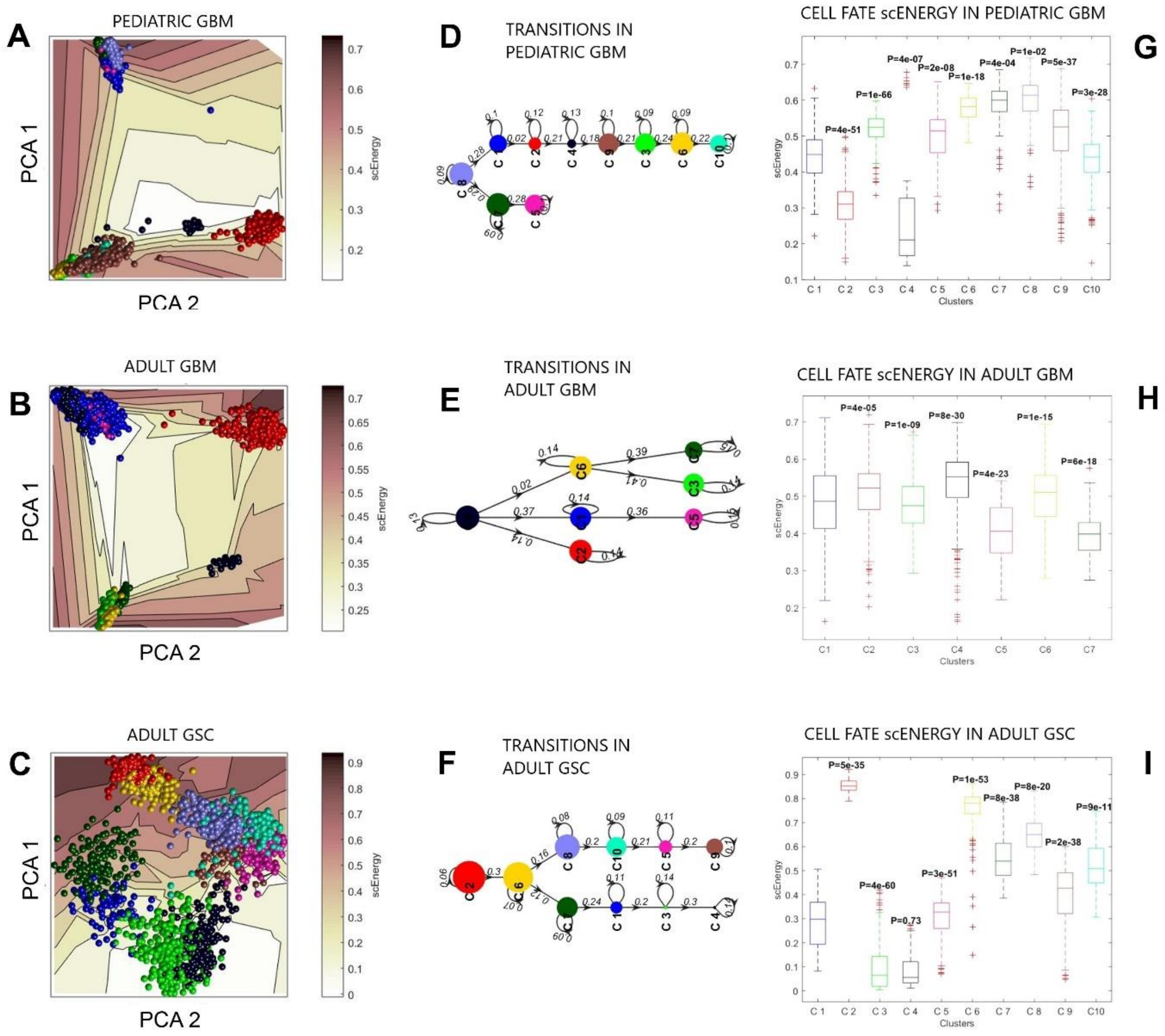


**Figure S2. Cell fate trajectories and cell-state transition probabilities inferred from GBM and GSC energy landscapes.** A) 2D energy contour of Pediatric GBM Waddington landscape. B) 2D energy contour of Adult GBM Waddington landscape. C) Energy contour of Adult GSC Waddington landscape. D) Cell fate bifurcations in Pediatric GBM. The transition probabilities between the distinct cell fates are shown by the numerical values above the nodes and node size corresponds to the single-cell energy on the landscape (scEnergy). E) Cell fate bifurcations in Adult GBM. F) Cell fate bifurcations in Adult GSC. G) Boxplot of scEnergy per cell fate cluster in Pediatric GBM. H) Boxplot of scEnergy per phenotypic cluster in Adult GBM. I) Boxplot of scEnergy per cell state cluster in Adult GSC**.**

**Putative Functional Relationships were identified by the Network Centrality Measures.**

As highlighted in Figure 5 in the Main Text, the maximal network centrality measures identified key drivers of the information flow across the inferred GBM/GSC regulatory networks. Literature analysis further confirmed the identified drivers play key roles in GBM/GSC cell fate dynamics. For instance, GATA2 expression is involved in embryonic development and the self-renewal/maintenance of stem cells (Rodrigues et al., 2012; Menendez-Gonzalez et al., 2019;

Wang et al., 2015). FOXG1 is a neurodevelopmental TF (Bulstrode et al., 2017). TAZ has been identified as a prognostic indicator of GBM in previous studies (Mani et al., 2008). SATB2, YY1, OLIG1/2, and SOX6 were other critical transcription factors (TFs) identified as critical regulators of information flow in the complex networks driving GBM/GSC cell fate decisions. MTSS1 is involved in inhibiting the EMT (epithelial-mesenchymal transition) switch, a key regulator of cell fate transition dynamics in cancer stem cell differentiation and metastatic invasion (Yu et al., 2020). MTSS1 acts as an intracellular tumor suppressor and may be related to tumor metastasis (Yu et al., 2020; Schemionek et al., 2015). For example, EMP1 has been reported as a regulator of glioma stemness and was shown to have a high eigenvector centrality in the BT127 GSC cells. S100B, one of markers identified in the differential expression analysis of GBM cells is a transcriptional target of SOX6, which was shown to have the highest betweenness centrality in adult GBM (Saito et al., 2007). The interconnectedness of the identified networks in their functional relationships from independent clustering algorithms further suggests our findings have clinically relevancy for potential targeted therapies.

Amongst all identified critical network dynamics regulators, ATL3 may be the most influential. It had the highest score for most of the centralities in all patient groups. ATL3 is a key regulator of ER (endoplasmic reticulum) network biogenesis (Lü et al., 2020) and may be a critical regulator for the interplay between gene expression and protein synthesis. However, our findings demonstrates its strong relevance to the information flow across the complex networks driving GBM cell fate dynamics.

**Estimates of algorithmic complexity provide key insights into genes differentiating phenotypes.**

The BDM method computes the K-complexity estimate in bits, of the gene expression counts of the PIDC network nodes with the highest PID scores (see Methods, Main Text). A block size of 12, and an alphabet size of 2 were used for the computations (Figure S3). Machine Learning (ML) classifiers, namely the AdaBoost Random Forest (RF) and Support Vector Machines (SVM) as described in the Supplementary Methods Section (See Below) were used in the study to determine which of these influential nodes of the network can distinguish the three distinct patient groups.

Notably, we observed a 1.00 (100%) classification accuracy with a 0.0 mean square error for the AdaBoost RF classifier on FOSB K-complexity scores (Fig S3A). Equivalent classification results were observed for a 0.2 validation size demonstrating the robustness of the classification on different test: train sizes (Fig S3A). The linear kernel SVM had a 90.909% classification accuracy on the FOSB K-complexity scores with a mean-square error of 0.0909 (data not shown). The cross-validation curve for the RF classification had 72.22 ± 20.79 % cross-validation (CV) accuracy, which was obtained for the 0.5 test size and 93.33 ± 9.43 % CV accuracy for the 0.2 test size (Fig S3B). Larger sample sizes would be needed to reduce the CV accuracy's uncertainty and improve the training of the classifier (Fig S3B).

The AdaBoost RF classifier's performance on HMGB1 K-complexity scores showed a 100% accuracy and 0.0 mean square error with both 0.5 test size and 0.2 test size (Fig S3C). Identical classification results were obtained for the SVM classifier on HMGB1 K-complexity scores. The f1 score of 1.00 in both classifiers indicate the binary classifiers can accurately distinguish the three patient groups using the K-complexity scores of the HMGB1 gene expression counts. A 100% CV accuracy is obtained for the RF classifier on the HMGB1 K-complexity scores (Fig S3D). As shown, the curve in turquoise denoting the cross-validation score exhibits an optimized training as the training set size increases and reaches the maximum training score of 1.00 accuracy (violet dashed line). The minimized grey shaded regions indicate the reduced uncertainty in the training performance and show that HMGB1 K-complexity is a robust marker distinguishing the three patient groups (Fig S3D). 100% classification accuracy was obtained with both a 0.5 test size and 0.2 test size on the EGR1 K-complexity scores using the linear kernel SVM (Fig S3E). The f1- score was found to be 1.00 (Fig S3E). The CV accuracy for the SVM's performance on EGR1 algorithmic complexity scores was found to be 80.56 ± 14.16% (Fig S3F).

There are nonetheless limitations to this analysis. For example, two classes (i.e., GBM and GSC) could have made the classification accuracy of other genes' K-complexity better. Alternately, sets of gene interactions could have been used for the K-complexity assessment. These analyses were not performed since it was demonstrated that all three genes identified by the ML classification on K-complexity estimates, namely, HMGB1, EGR1, and FOSB are directly interlinked by YY1, one of the critical transcription factors identified in our network science measures (See Main Text). Further, K-complexity was used to identify robust discriminants of the three patient-groups amidst the regulatory network markers. However, the OACC had a network perturbation analysis feature exclusively for unweighted networks which remains to be epxloited for weighted complex networks. Shannon entropy was discarded as an information theoretic in assessing the signaling complexity as it was roughly the same value of 1.2-1.3 for all nodes and hence, was verified to be a poor measure of signaling complexity as suggested by Zenil et al. (Zenil et al., 2019). However, other algorithmic information dynamics (AID) tools including compression algorithms (LZW, Hauffman coding, RLE, etc.) and a 2D-Block Decomposition Method (BDM) could also be deployed to study the graph complexity of the recoconstructed regulatory networks with perturbation analysis (Zenil et al., 2019).

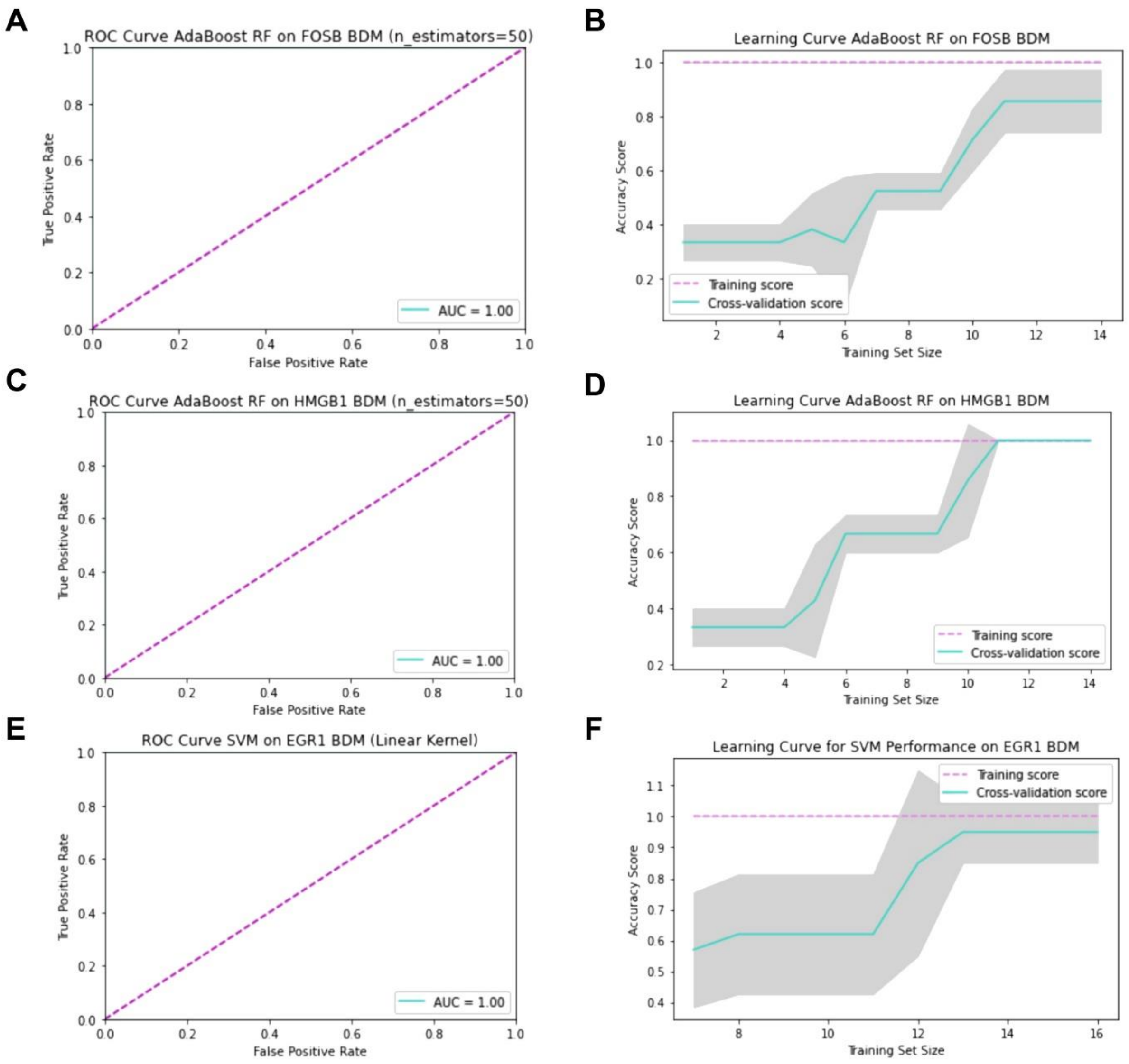


**Figure S3. Machine Learning Classifiers identify key genes capturing the algorithmic complexity of GBM networks.** The AdaBoost random forest (RF) and support vector machine classifiers were assessed on the K-complexity BDM scores of the genes with the top PID scores on the inferred PIDC networks. The AdaBoost RF classifier was kept at default set of hyperparameters: a learning rate of 1.0, and 50 n_estimators, while the SVM had a learning rate C= 1.0. The receiver operating characteristic (ROC) curve provides a visual representation of the classification algorithm's accuracy. Farther the turquoise area under curve (AUC) line is from the dashed violet curve at 45 degrees, the higher the classification accuracy. A) AdaBoost_RF classifier performance on FOSB BDM scores. B) AdaBoost_RF cross-validation learning curve on FOSB (0.5 validation size). C) AdaBoost_RF classifier performance on HMGB1 BDM scores. D) AdaBoost_RF cross-validation curve on HMGB1 BDM scores. E) SVM classifier performance on EGR1 BDM scores. F) SVM classifier cross-validation curve on EGR1 BDM scores.

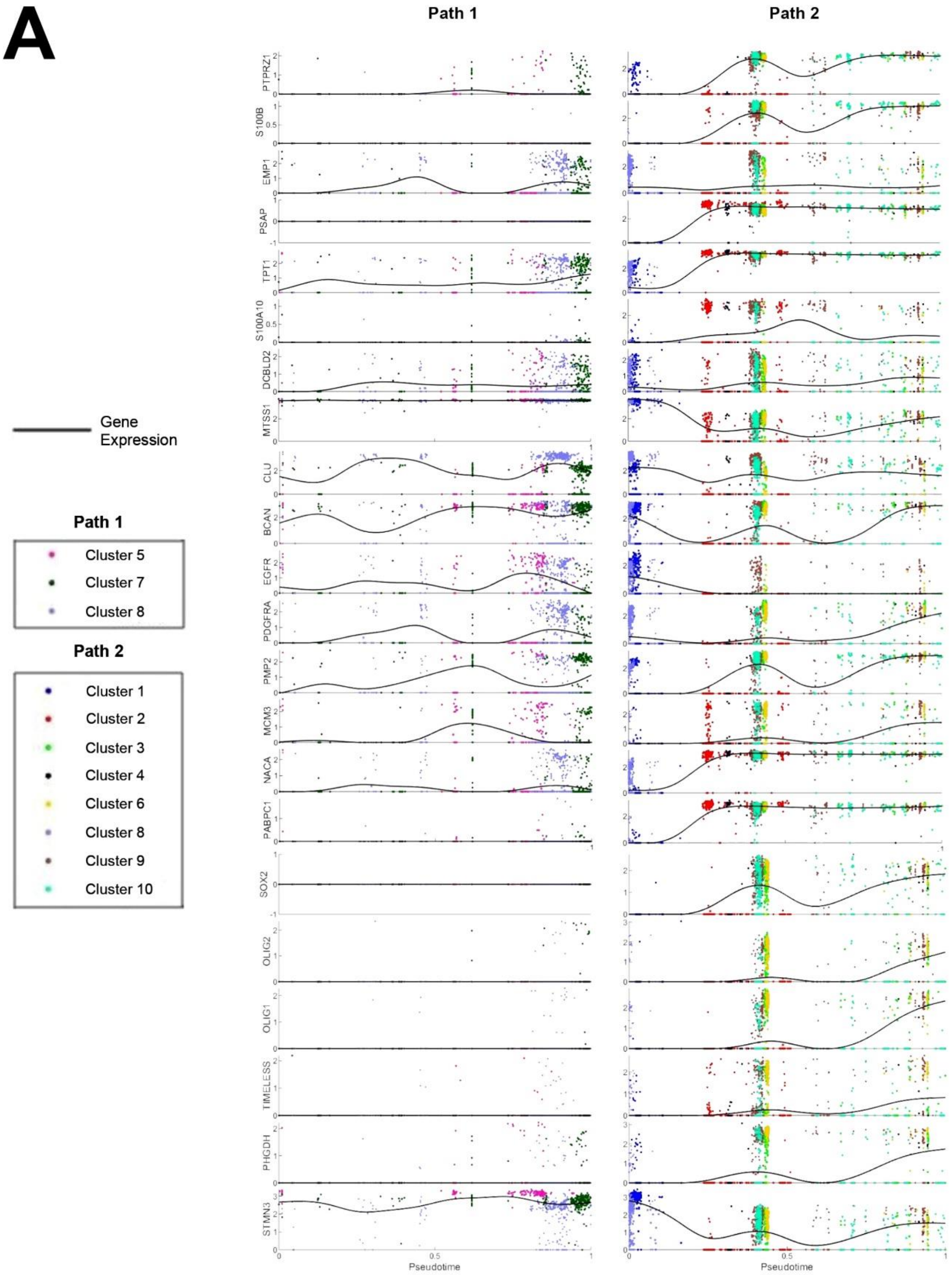
A
Path 1
Path 2
PTPRZ1
S100B
EMP1
PSAP
TPT1
S100A10
DCBLD2
MTSS1
CLU
BCAN
EGFR
PDGFRA
PMP2
MCM3
NACA
PABPC1
SOX2
OLIG2
OLIG1
TIMELESS
PHGDH
STMN3
Pseudotime
Pseudotime
Gene Expression
Path 1
Cluster 5
Cluster 7
Cluster 8
Path 2
Cluster 1
Cluster 2
Cluster 3
Cluster 4
Cluster 6
Cluster 8
Cluster 9
Cluster 10

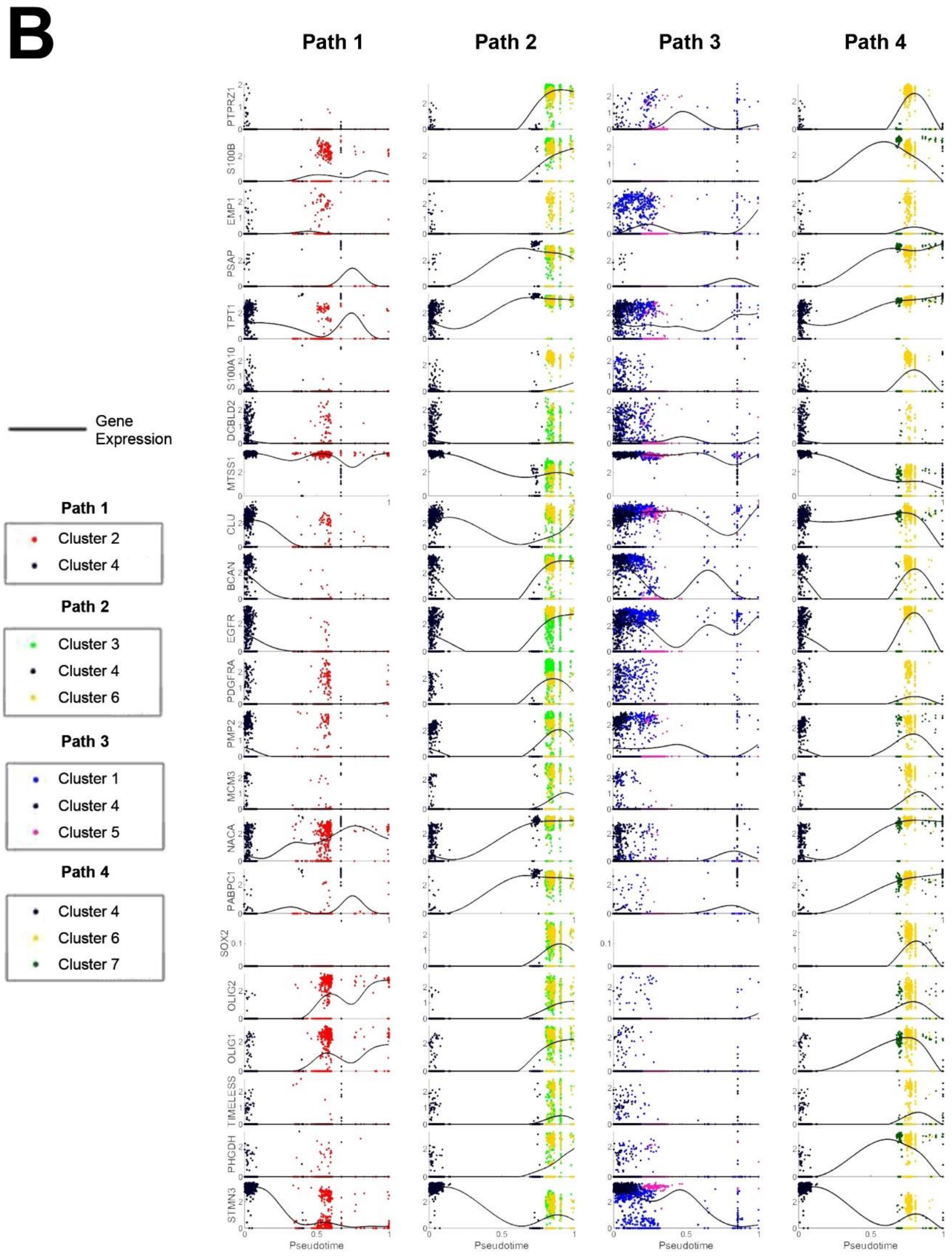
B
Path 1
Path 2
Path 3
Path 4
PTPRZ1
S100B
EMP1
PSAP
TPT1
S100A10
DCBLD2
MTSS1
CLU
BCAN
EGFR
PDGFRA
PMP2
MCM3
NACA
PABPC1
SOX2
OLIG2
OLIG1
TIMELESS
PHGDH
STMN3
Pseudotime
Gene Expression
Path 1
Cluster 2
Cluster 4
Path 2
Cluster 3
Cluster 4
Cluster 6
Path 3
Cluster 1
Cluster 4
Cluster 5
Path 4
Cluster 4
Cluster 6
Cluster 7

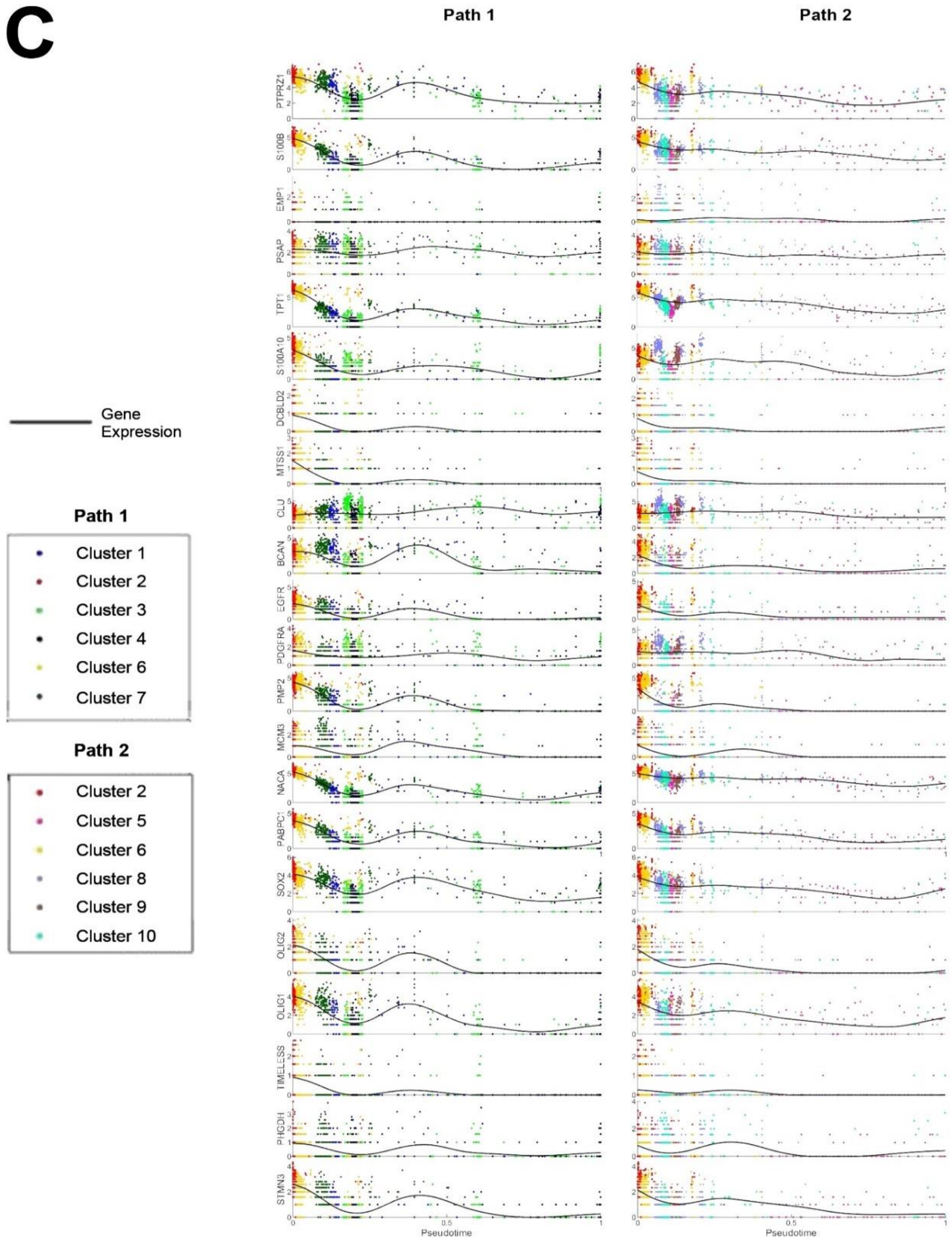


**Figure S4. Pseudotime dynamics of cubic-spline smoothed average, normalized gene expression patterns in cell fate differentiation (extension of Figure 5).** A) Pediatric GBM, B) Adult GSC, and C) Adult GSC. Oscillatory dynamics are observed in temporal expression dynamics during cell fate transitions indicating the possibility of complex attractors steering GBM/GSC cell fate decisions.

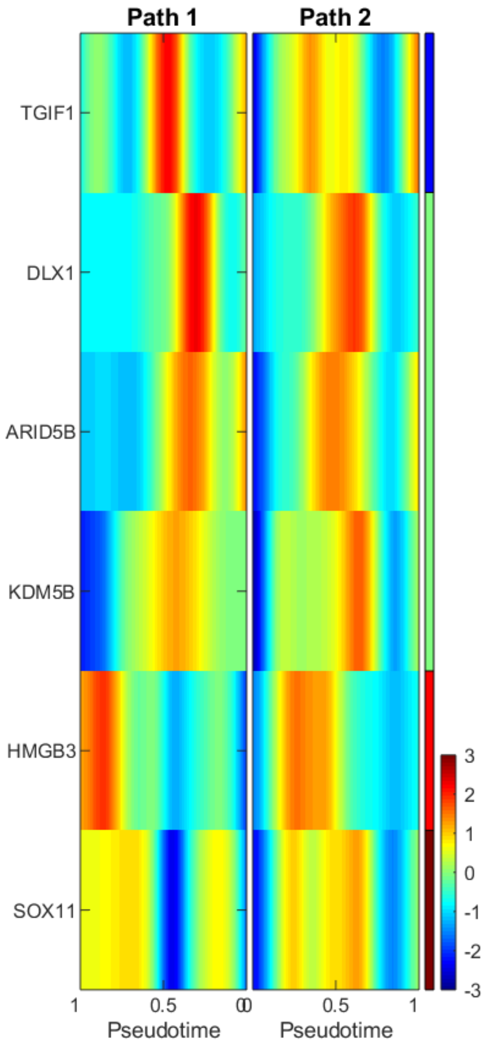


**Figure S5. scEpath Cell Fate Transition Heatmap for Pediatric GBM (n=8, N=1943 cells).** The transition markers controlling pseudotemporal differentiation dynamics when all eight pediatric GBM samples from Neftel et al. (2019), removed in our quality control step, were included in trajectory inference analysis. Two developmental trajectories (transition paths) are observed. However, the list of transition markers has reduced. The SOX11 marker is a neuronal differentiation regulator, also observed in the adult GSC heatmaps in Figure 1F. The color bar denotes the normalized gene expression.

**Biological significance of identified genes and transcription factors**

Previous studies have shown that EMP1 facilitates the aggressivity and metastatic potential of GBM cells potentially by activation of the PI3K/AKT/mTOR signaling pathway (Miao et al., 2019). MTSS1 is a metastasis suppressor gene, the methylation profile of which has been shown to steer GBM migration (Luxen et al., 2017). MTSS1 is also involved in controlling epithelial-mesenchymal transition (EMT) and hence, regulating GSC stemness in GBM populations (Yu et al., 2020). OLIG1/2 are master neurodevelopmental transcription factors essential for GBM propagation and GSC stemness (Suvà et al., 2014). PTPRZ1 encodes a member of the receptor protein tyrosine phosphatase involved in developmental processes of the central nervous system (Harroch et al., 2002). Studies have shown PTPRZ1 promotes GBM invasion, coordinates GBM cellular homeostasis, and regulates the cross- talk between tumor-associated macrophages (TAM) and GSC cells (Shi et al., 2017, Bhaduri et al., 2020). S100B encodes a glial-specific calcium-binding protein involved in complex processes such as cell cycle progression and astrocyte differentiation (Brozzi et al., 2009). Interestingly, S100B also has been shown to regulate glioma growth via TAM-associated chemoattraction (Wang et al., 2013). Our findings thus suggest a possible regulatory feedback loop between PTPRZ1 and S100B in GBM cell fate dynamics.

The abnormally high PIDC scores for the TIMELESS and PHGDH interaction observed in the BT127 adult GSC sample suggests a possible mechanism distinguishing GSC from GBM. PHGDH is a key enzyme in metabolic energy transduction that is also involved in chemoresistance and promoting glycolytic GBM phenotypes (tumor metabolism regulation) (Ehmsen et al., 2013, Engel et al., 2020). Given that PHGDH is expressed highly in developing glial and astrocytic cells (Yamasaki et al., 2001), this interaction may be the primary source of energy used by GSCs, namely, oxyglutarate or pyruvate derivatives, as predicted by the Warburg effect (Liu et al., 2013).

Via its interaction with EGFR, TAZ promotes cancer stemness and phenotypic plasticity (Gao et al., 2021) and phenotypic plasticity and differentiation to mesenchymal phenotypes via the EMT pathway (Bhat et al., 2011), hence GBM cells expressing TAZ may acquire GSC properties and behaviors (Mani et al., 2008). TAZ may therefore play a critical role in GBM therapeutic resistance, and further promotes hybrid OXPHOS/glycolytic phenotypes providing a metabolic advantage to adapt to the fluctuant microenvironment (Jia et al., 2017). GATA2 is a hematopoietic transcription factor essential to the maintenance and differentiation of hematopoietic stem cells that has been found to associated to hematopoietic malignancies (Menendez-Gonzalez et al., 2019, Rodrigues et al., 2012) and promote glioma progression through the EGFR/ERK pathway (Wang et al., 2012). MECOM is an oncogenic TF known for its downstream signaling with TGF-$\beta$ and cell cycle regulators that is associated with poor GBM prognosis (Hou et al., 2016). FOXG1 is a neurodevelopmental TF highly expressed in GBM cells which, together with elevated expressions of SOX2, promotes GSC phenotypes in GBM (Bulstrode et al., 2017). SATB2 was recently shown as a driver of GBM progression preferentially expressed by GSCs (Tao et al., 2020) and may be a critical gatekeeper of cancer stemness in GBM cell populations (Roy et al., 2020). Hence, it should be further investigated as a clinical drug target.

The overexpression of the transcription factor YY1 interacts with NF-kB dependent pro-inflammatory signaling pathways in GBM-immune interactions (Polisetty et al., 2012) and has previously been reported in GBM and shown to promote chemoresistance (Baritaki et al., 2009). SOX6 has functional relevance in the normal development of the central nervous system (Azim et al., 2009) and the role of SOX family members of TFs in cancers and maintenance/control of pluripotent stem cell fates is well-established (Grimm et al., 2020; Hagiwara, 2011; de la Rocha et al., 2014). S100B, one of the markers identified in our differential expression analysis of GBM cells, is a transcriptional target of SOX6 (Saito et al., 2007). TPT1 is a cancer-associated gene that encodes a protein known to promote glioma growth and progression, primarily via calcium binding and microtubule-cytoskeletal dynamics (Gu et al., 2014). Further, TPT1 was established as essential for activating pluripotency (stemness) genes such as Oct-4 and Nanog upon nuclear transfer (Koziol et al., 2007). Previous findings have shown TPT1 is key to phenotypic reprogramming in cancers and hence, regulates complex adaptive processes such as tumor reversion (Amson et al., 2013). Hence, our findings suggest TPT1 promotes GSC stemness. Interestingly, ATL3, whose function remains poorly investigated, occupied the highest number of maximal centrality measures across all patient groups. Given that many of the identified differentially expressed signals in our GBM networks are involved in calcium signaling, we speculate that ATL3 may serve as a hub for regulating ER stress and extracellular vesicles transport mediating GBM cell-cell communication networks (Polisetty et al., 2012).

PRC2 dynamics are critical for the regulation of developmental-associated genes and control the epigenetic plasticity of pediatric gliomas. PRC2 is vastly inhibited or dysregulated in pediatric gliomas, especially the oncohistone variant H3K27M (Schwartzentruber et al., 2012), and was verified to interact with all six transition genes as shown in Table 3. We believe PRC2-YY1 interactions may also be controlling the expression dynamics of other key TFs identified in our analyses. For example, YY1 and EZH2 are known to control cell cycle regulation and transcriptional dynamics by interaction with TAZ (Hoxha et al., 2020). As such, we speculate dysregulated or impaired YY1 expression may allow GBM to overcome transcriptional repression of critical cell cycle control genes and thereby promote gliomagenesis and GBM progression. On the other hand, ATF3 is known for encoding CREB protein family TFs that are involved in calcium-mediated complex cellular processes regulating cancer-immune dynamics (Thompson et al., 2009). The roles of other high importance TFs identified in the network analyses such as FOXG1, SATB2, and MECOM with respect to the above-listed genes/TFs remains unelucidated in cancer/GBM dynamics.

### SUPPLEMENTARY METHODS

#### Single-cell Data Analysis

Smart-seq2 whole transcriptome amplification, library construction, and sequencing were taken from Filbinet al., 2018, Picelli et al., 2014, Tirosh et al., 2016b, and Venteicher et al., 2017 (Neftel et al., 2019). For a subset of samples in (Neftel et al., 2019), single cells were processed via the 10X Chromium 30 Single Cell Platform using the Chromium Single Cell 30 Library, Gel Bead and Chip Kits (10X Genomics, Pleasanton, CA). 7,000 cells were added to each channel of a chip partitioned into Gel Beads in Emulsion (GEMs), followed by cell lysis and barcoded reverse transcription of RNA in droplets-Seq. De-emulsion was followed by amplification, fragmentation, and addition of adaptor and sample index (Neftel et al., 2019). Similar treatment conditions were applied for the GSC count matrices with > 69,000 adult GSC cells extracted from 26 patients (Richards et al., 2021).

Among filtered cells, an average of 5,730 genes per cell were found as a quality measure. Expression levels were quantified as $E_{i,j} = log2\left(\frac{TPM_{i,j}}{10} + 1\right)$, where $TPM_{i,j}$ refers to transcript-per-million for gene $i$ in sample $j$, as calculated by RSEM (Neftel et al., 2019). TPM values were divided by 10 given that the complexity of single cell libraries was estimated to be on the order of 100,000 transcripts. For the remaining cells, the aggregate expression of each gene was calculated as $Ea_i = log2(average(TPM_i) + 1)$, for $i = 1 \dots n$. (Neftel et al., 2019) then defined the relative expression over the remaining cells by centering the expression levels per gene, i.e., $Er_{i,j} = E_{i,j} - average[E_i]$.

#### scEpath Waddington Landscape Reconstruction Algorithm

The scEpath algorithm performs PCA analysis on the energy matrix $E = (E_{ij})$ and fits a potential energy surface using piecewise linear interpolation over the first two PCA components and the single cell energy (scEnergy) of each cell. Cells are then colored according to unsupervised clustering which groups cells with similar gene expression patterns (transcriptional states). The energy of each cell state (scEnergy), $E_j$, on the Waddington landscape is computed according to:

$$E_j(y) = \sum_{i=1}^{n} E_{ij}(y) = -\sum_{i=1}^{n} y_{ij} ln \frac{y_{ij}}{\sum_{k\in N_i} y_{kj}},$$

where $y_{ij}$ represents the normalized gene expression level (between 0 and 1) of gene $i$ and cell $j$, and $N(i)$ is the neighborhood of node-$i$ in the network. Each gene is assigned a local energy state $E_{ij}$ (Jin et al., 2018). The scEnergy is combined with a distance-based measure and structural clustering to reconstruct the 3D energy landscapes.

The cell-state on the scEpath energy landscape corresponds to which discrete bin its mRNA levels fluctuate within (Jin et al., 2018). The cell states distribution on the scEnergy landscape can thus be defined as attractors, a term from dynamical systems theory used to describe a causal pattern to which the cell state dynamics are bound. Assessing the fractal dimension of this attractor (cell state patterns) for each patient group's energy landscape provides key insights into the complexity of the cell states and the transition gene dynamics governing their cellular decision-making.

**Mapping Pseudotemporal Ordering and Cell Lineage Bifurcations in GBM/GSC Cell Fates**

scEpath performs Principal Component Analysis (PCA) on the energy matrix and uses the first two PCA components as the reaction coordinates followed by structural clustering of cells via an unsupervised cell-cell similarity metric called single-cell interpretation via multikernel learning (SIMLR) (Jin et al., 2018). To infer cell lineages, scEpath constructs a probabilistic directed graph in which nodes represent phenotypic clusters (attractors), with edges weighted by cell state transition probabilities (Figure S2). By default, scEpath defines the cell clustering patterns as the set of cells occupying 80% percent of the total energy in each cluster. The cell state transition probability such that a cell state will be in cell cluster *k* with a particular scEnergy was calculated using the Boltzmann–Gibbs distribution by the scEpath algorithm (Jin et al., 2018). The directions of the probability flow are determined by the energy flow with significant changes from high to low. scEpath learns the maximum probability flow in the probabilistic directed graph defined by a weighted matrix $W$ determined by the gene expression counts. This problem is equivalent to finding the minimum directed spanning tree by setting the edge weights to be $1 - W$. Thus, scEpath implements Edmonds' algorithm to determine the minimum directed spanning tree (MDST) connecting the cell clusters (attractors) and hence, the candidate cell lineage bifurcations of cell fate decisions (Jin et al., 2018).

**PIDC Network Inference Algorithm**

We used *Julia LightGraphs* to infer the PIDC network. This algorithm works by constructing undirected simple weighted graphs that optimize the shortest path of vertices/nodes based on the weights of the network (i.e., the PID scores). The network nodes are ordered by the top PID scores in decreasing order. Given three variables (genes) $X$, $Y$, and $Z$ on the network, partial information decomposition (PID) maps the information obtained from a source set of genes $S$={$X, Y$} about the target gene $Z$. The information is redundant, synergistic, and unique. PID is an information-theoretic similar to pairwise mutual information, taking into consideration the information dynamics between three-variables at a time (instead of pairwise-correlations) (Chan et al., 2017). Thus, the PID score between the source gene set $S$ and the target gene $Z$ is given by $I(X; X, Y)$ :

$$I(X;X,Y) = Synergy(Z;X,Y) + UniqueY(Z;X) + UniqueX(Z;Y) + Redundancy(Z;X,Y),$$

where $Unique\ (Z;X)$ is the unique information between the source gene $X$ and target gene $Z$ when the other source gene is $Y$. The PIDC inference algorithm calculates the PID scores.

The PIDC inference algorithm may be simply defined as follows. PID values are estimated for every gene triplet (with each gene treated as the target gene in the others), and from these the proportional unique contribution (PUC), represented as $U_{X,Y}$ (as defined below), is estimated for every pair of genes. For each gene $X$, an empirical distribution $f_X(u)$ is estimated. The confidence of an edge between a pair of genes depends on the corresponding cumulative distribution functions $F_X(u)$(assumed as either a Gamma or Gaussian empirical probability distribution, for each gene within the pair). These confidence scores are then used to rank all possible network edges. The *Julia LightGraphs* algorithm uses the cumulative probability distributions for each gene to obtain a final confidence score for network edges (Chan et al., 2017). The corresponding PID scores are obtained as output of the algorithm.

We define the PUC between two genes X and Y as the sum of this ratio calculated using every other gene $Z$ in a network:

$$U_{X,Y} = \sum_{Z \in S(x,y\}} \frac{Unique_z(X;Y)}{I(X:Y)} + \sum_{Z \in S(x,y\}} Unique_z(X;Y)$$

This measure captures the mean proportion of mutual information (MI) between two genes $X$ and $Y$.  Using an unsupervised Louvain community detection algorithm, the PID network is then inferred from the PID scores of the genes. Such network analyses coupled to the scEpath algorithm helped us identify gene modules which may exhibit oscillatory dynamics in gene expression as cells under-go state transition, and putative gene interactions which may be involved in regulating GBM/GSC cell fate choices (see Main Text).

**Machine learning and Block Decomposition Analysis.**

Binary classification was performed using Google Colab using Scikit-learn on the BDM scores of three classification groups (pediatric GBM, adult GBM, and GSC). The linear support vector machine (SVM) and AdaBoost random forest (RF) classifier modules (with hyperparameter tuning to optimal learning rates) from Scikit-learn were trained using both an 80:20 and a 50:50 training: testing size split with the BDM scores of the gene markers with the top PID scores identified in the PIDC networks for both the GRNs and TF networks. Seven patient samples were selected for each of the three groups for the classification training and validation, as defined above. GraphPad Prism 8.4.3 was used for additional statistical analyses.
